# Reshaping the Landscape of Locoregional Treatments for Breast Cancer Liver Metastases: A novel, intratumoral, p21-targeted percutaneous therapy increases survival in BALB/c mice inoculated with 4T1 triple negative breast cancer cells in the liver

**DOI:** 10.1101/2024.11.27.625706

**Authors:** Bryan S. Margulies, Morakot Likhitpanichkul, Debu Tripathy

## Abstract

Patients with disseminated metastatic disease from breast cancer are likely to have liver involvement in >50% of cases at some point during disease progression. These patients have a poor prognosis; and, when treated with the standard of care systemic therapy they have a median survival of <9-months. Increasing survival in breast cancer patients will likely require the administration of better therapies that are specifically targeted to treat distant metastases. One approach to increasing treatment efficacy for breast cancer liver metastases is through the application locoregional therapies. Locoregional therapies are an appealing interventional approach for breast cancer patients with liver metastases since these tumor lesions are accessible via minimally invasive procedures that can be administered using either ultrasound or CT imaging. Current locoregional therapies to treat breast cancer liver metastases are non-specific and have not produced significant increases in survival. The goal of this study was to design and test a targeted locoregional therapeutic intervention for breast cancer liver metastases. The lead candidate, a fixed-dose small-molecule drug called MBC-005, was tested *in vitro* and then the efficacy was evaluated in a BALB/c mouse liver metastases model. A novel formulation of N-allyl noroxymorphone hydrochloride incorporated into an alginate-based gel overcomes many of the limitations associated with the administration of small-molecule drugs, which include solubility, off-target toxicity, and enzymatic degradation. *In vitro* results demonstrated that MBC-005 mediated its anti-tumorigenic effect through a p21-dependent mechanism via a novel molecular pathway, in which N-allyl noroxymorphone component of MBC-005 stimulated the opioid growth factor receptor to increase p21 expression. Intratumoral administration of MBC-005 increased survival 3.9-fold in mice and significantly decreased tumor volume 4-fold. While many cytotoxic therapies increase p21 expression as a response to DNA damage, MBC-005 increased p21 expression independent cytotoxic DNA damage. MBC-005 did not induce off-target toxicity; and, as such, would be amenable to multiple rounds of administration. Nevertheless, it is notable that the positive effects of MBC-005 treatment on increasing survival and decreasing tumor volume in mice was achieved using a single dose.

## Introduction

The delivery of small-molecule therapies to treat cancer has long been complicated by chemical properties that are incompatible with organic systems, which includes hydrophobicity, solubility, enzymatic degradation, off-target toxicity, and renal clearance [1–4]. Nevertheless, patients would certainly benefit from novel, targeted small-molecule therapies if these delivery challenges could be addressed. Recently, we developed a formulation of N-allyl noroxymorphone hydrochloride that overcomes several of these hurdles, specifically solubility, off-target toxicity, and enzymatic degradation. N-allyl noroxymorphone is slightly hydrophobic when pH >7, which limits the ability to achieve high concentrations in solution [5, 6]. Initial in vitro studies indicated that concentrations of N-allyl noroxymorphone that are therapeutically useful would be greater than could be achieved using this initial formulation. As such, we determined that N-allyl noroxymorphone would need to be incorporated into a delivery system that could overcome its hydrophobicity.

In our initial studies we found that a novel formulation of N-allyl noroxymorphone (BC-003) stopped cell proliferation through the upregulation of p21 [7]. The p21 transcription factor is a master regulator of cell proliferation and differentiation [8, 9]. Given the role p21 plays in regulating cell proliferation, it is a potential target for treating cancer [10]. Initial in vitro studies in breast cancer tumor cells suggested that N-allyl noroxymorphone could be an effective therapy. This result led us to the present investigation, in which we developed a novel delivery system for our novel formulation of N-allyl noroxymorphone to overcome issues surrounding hydrophobicity so that a therapeutically useful concentration could be achieved. This new formulation of N-allyl noroxymorphone, MBC-005, was then tested in mouse livers inoculated with breast cancer cells that were subsequently treated with the MBC-005 formulation.

Unlike other locations in which breast cancer metastases are common, liver metastases are amenable to the local administration of therapy. Additionally, patients with disseminated metastatic disease are likely to have liver involvement in 50% to 62% of cases at some point during their disease progression [11, 12]. The current standard of care (SOC) for patients with breast cancer liver metastases is systemic chemotherapy and locoregional intervention if their liver metastases are symptomatic [13]. However, patients with liver metastases have a poor prognosis, with the median survival rate being <9-months, when receiving conventional SOC systemic chemotherapy for their disease [11, 14–17]. In addition, patients with liver metastases often experience serious complications that arise from the presence of the metastases that include rapidly progressive hepatic failure, refractory ascites, portal vein thrombosis, and nutritional compromise [18].

Patients with liver metastases that are amenable to resection are in the minority, with only 5% to 25% being suitable candidates for hepatectomy [19]. Patients with unresectable metastases in the liver are treated using one of the following local interventions [20, 21]: trans arterial chemotherapy embolization (TACE), radio-frequency embolization (TARE) / selective internal radiation therapy (SIRT), thermal ablation, cryotherapy ablation, chemical ablation, and microwave ablation. Ablative minimally invasive interventions are considered better alternatives for use in palliative care and are the only option for unresectable metastatic disease; however, they are being used more often as adjuncts to surgical resection [20, 21]. TACE and TARE are the preferred therapeutic intervention to treat multifocal, unresectable, liver metastases from breast cancer [11, 22–24]. However, TACE and TARE both have significant complications that include hepatic failure, biliary obstruction, occlusion of the end-arterial supply to the biliary ducts that results in ischemic necrosis, biloma formation, the absence of a functioning sphincter of Oddi that compromises the sterility of the biliary tree and elevates the risk of liver abscesses [11, 22–24]. Further, TARE and TACE only increase 1-year survival approximately 30% compared to patients who are administered SOC chemotherapy [11, 22–24].

One of the limitations in evaluating locoregional therapies is that there are very few randomized clinical trials. Recent studies, such as NRG-BR002, failed to show a progression-free survival benefit from local stereotactic X-radiation therapy. However, despite this disappointing result, patients with tumors that are phenotypically homogenous have benefitted from stereotactic X-radiation, such as in the SABR-COMET study. Part of our motivation for the current work rests in the belief that locally administered targeted biologic therapies have yet to be rigorously tested.

The incidence of breast cancer continues to rise, with 367,220 estimated new diagnoses in 2024, despite the improvement in overall survival for breast cancer patients during last several decades [25]. This increase in the incidence of breast cancer is correlated, in part, with an increase in the aging population, with 57% of diagnoses occurring in people ≥65-years of age [26]. This is of special concern since this population is expected to grow to 81 million in 2040 [26]. Further, in 2022, 4.1 million people were living with breast cancer, 4% (164,000) of whom have distant metastases [25], a number that is also expected to grow with the corresponding increase in survival. A very large number of breast cancer survivors are at risk for relapse or recurrence. Relapse and recurrence, depending on the breast cancer phenotype, can range from 5.8% to 22.8% for non-triple negative breast cancer (TNBC), 40% for TNBC, and 50% for inflammatory breast cancer [27]. In patients experiencing relapse or recurrence, the incidence of distant metastases is high [28]. Ultimately, in 2024 the numbers of estimated deaths are expected to be 42,250 and 5-year survival is expected to remain at ≤31% for stage IV disease [25].

Increasing survival in breast cancer patients overall will likely require the administration of better therapies that are targeted specifically for patients with distant metastases. MBC-005 was designed to fulfill this need. The goal of this study was to design and test a locoregional therapeutic intervention to treat breast cancer liver metastases by optimizing the material properties of MBC-005, testing MBC-005 *in vitro*, and then evaluating the efficacy of MBC-005 in a BALB/c mouse liver metastases model. We hypothesized that an optimally designed MBC-005 would inhibit breast cancer growth in vitro and in vivo. Further, we hypothesize that MBC-005 will be effective through induction of p21, a known tumor suppressor gene, as its primary mechanism of action.

## Results

### Design of MBC-005

The following key parameters were identified for MBC-005: it must be flowable through a syringe, viewable under ultrasound (i.e., echogenic), and have an elution rate for N-allyl noroxymorphone of ≤0.2-mg/hr^1/2^. A sodium alginate/gum Arabic mixture (1:4.43 w/w) was empirically found to possess the handling properties needed to be injectable through a syringe when a lanthanum carbonate/iron (III) sulfate hydrate catalyst (1:14 w/w) was used to initiate hydrogel curing. Viscometry was used to estimate the injectability of MBC-005 using a tuning-fork viscometer. Time dependent changes in viscosity were observed, with the maximum viscosity of 10,163.8-mPa•s (N=8; SD ±1683.4) was reached at 560-sec (i.e., 9-minutes and 20-sec) over the 1800-sec (i.e., 30-minutes) experimental period (**Figure 1**). The range observed for the viscosity values likely relate to differences in humidity, which ranged from 14.2% to 30.9% during the experiments, while the mean temperature was 19.68° C varied less than 1° C (SD ±0.396° C). Interestingly, the material exhibited endothermic behavior during testing, with the average temperature decreasing 1° C for the material.

**Figure 1:**
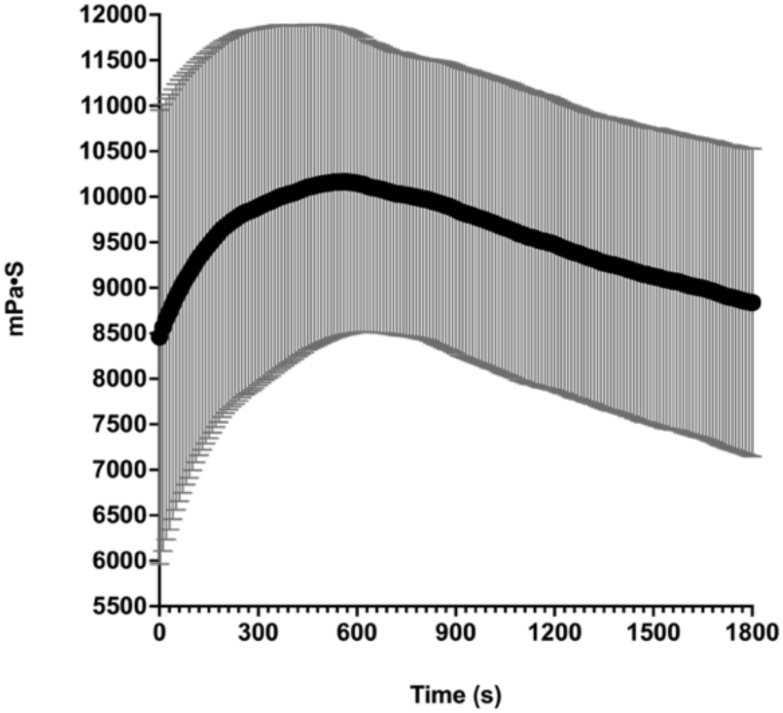
Viscosity assessed using a tuning-fork viscometer. Experiments were run for 1800-sec (i.e., 30-minutes) on N=8 samples. Humidity ranges from 14.2% to 30.9% for the experiments. The ambient temperature ranged from 18.9° C to 20° C. The average maximum viscosity was 10,163.8-mPa•s (SD ±1683.4-mPa•s) measured at 560-sec.

Previous work had shown that the average maximum force achievable during syringe injection is 79.8-N [29] while Vo et al. demonstrated that 4.9-N to 23.3-N of force was needed to inject water using a 5-mL syringe with a 25-G needle. Based on the viscosity data, we would expect that MBC-005 would require 10.163-N of force for injection, which is well below the 23.3-N force identified by Vo et al. and aligns well with our experience. However, we did find that a 16-G to 21-G needle was needed for a controlled injection of approximately 0.1-cc/s, with this range of needle sizes and injection rate consistent with clinical practice [30, 31]. The ability to inject MBC-005 using a syringe with an 18-G needle was further confirmed by demonstrating that it could be injected into bovine livers, which was imaged using ultrasound guidance (**Figure 2**).

**Figure 2:**
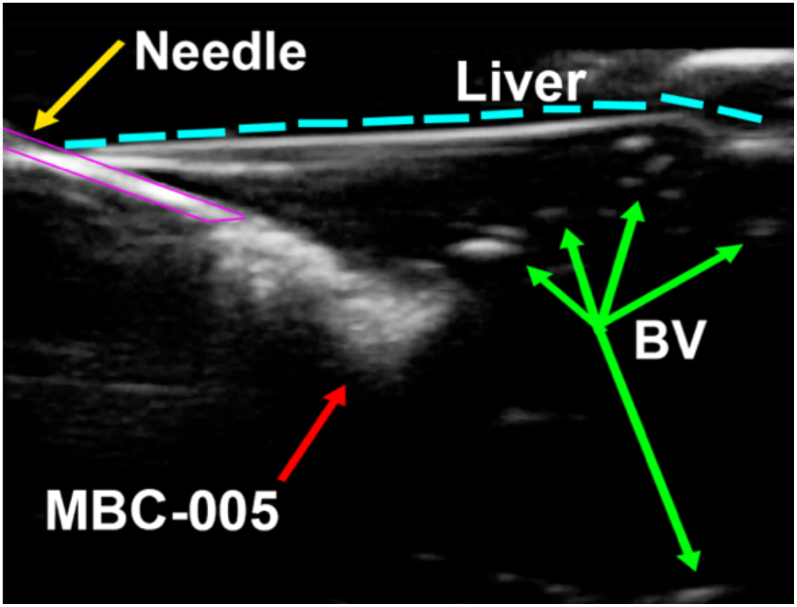
Echogenic properties were assessed using a portable ultrasound, similar to the devices that are used clinically. MBC-005 was loaded (3-mL) into a 5-mL syringe with an 18-G needle. Samples were injected into bovine liver samples embedded in glycerol.

The elution rate of the N-allyl noroxymorphone was empirically determined for the MBC-005 formulation using the Hixson-Crowell cube-root law for diffusion from a hydrogel, which models Higuchi diffusion kinetics using the following equation: 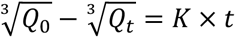. In this equation, *‘Q_0_’* is the starting amount of N-allyl noroxymorphone, ‘*Q_t_*’ is the amount of N-allyl noroxymorphone at time ‘*t*’, ‘*K’* is the rate constant, and ‘*t*’ is time. ‘*K*’ is a dimensionless constate that is derived empirically from the material properties of a material; and, as such ‘*K’* is an intrinsic property of MBC-005. Data were collected hourly using UPLC from which the quantity of N-allyl noroxymorphone was measured. A best-fit line modeled on the Hixson-Crowell equation was then computed (**Figure 3A**) for two different samples with an R^2^ of 0.928 and 0.8468, respectively. The mean release rate was 0.1496-mg/hr^1/2^.

**Figure 3:**
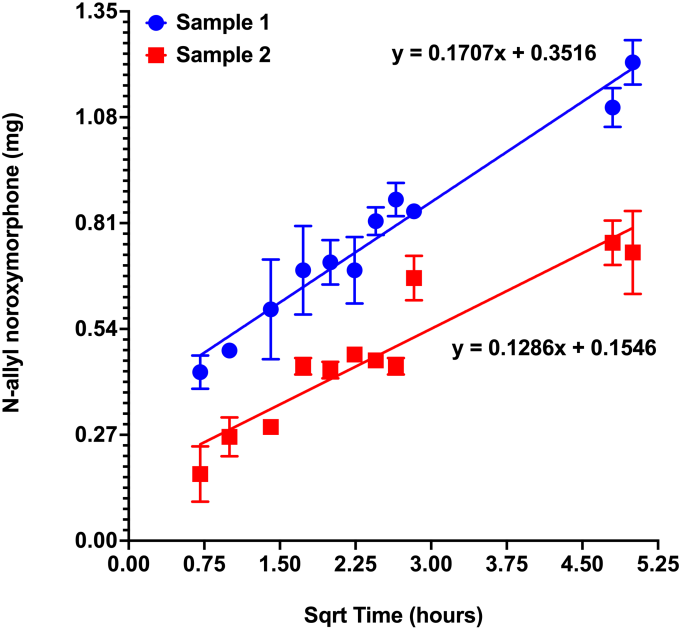
**(A)** Release rate (i.e., elution) of N-allyl noroxymorphone hydrochloride evaluated using UPLC from two samples of MBC-005. The equations predict that at 24-hours there will be between 0.798-mg and 1.205-mg of N-allyl noroxymorphone hydrochloride remaining.

A Design of Experiments approach was used to create a design space (e.g., the mathematical model), which was generated using a D-optimal quadratic model. D-optimal design uses an iterative algorithm that minimizes the parameter covariance. The optimal MBC-005 composition (**Figure 3B**) was identified using this model based on maintaining the lanthanum concentration above 25% and the elution rate ≤0.2-mg/hr^1/2^. The result of this analysis was the identification of an optimal MBC-005 formulation that is composed of 53% sodium alginate (w/w), 12% gum Arabic (w/w), and 26% lanthanum carbonate (w/w). This optimal MBC-005 formulation has an elution rate of 0.14647-mg/hr^1/2^.

**Figure 3:**
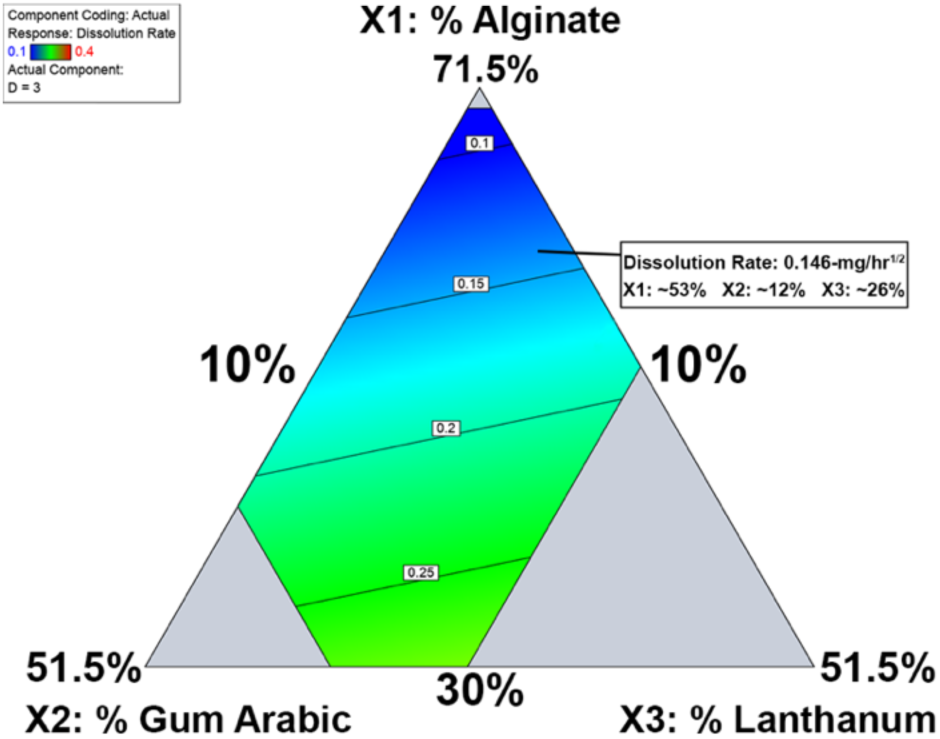
**(B)** Contour plot showing N-allyl noroxymorphone hydrochloride elution using different compositions of MBC-005. The contour plot generated predicts an optimal elution rate at 24-hours of 0.14647-mg/hr^1/2^.

### MBC-005 Effects on Tumor Cytotoxicity

*In vitro* culture studies demonstrated that the small molecule N-allyl noroxymorphone reduced human breast tumor cell number in a dose-dependent manner 72-hours after administration for both the BT474 breast cancer cells (hormone receptor positive (HR+) / HER2+) and the HR-positive/HER2-negative MCF7 breast cancer cells when compared to the HUMEC primary breast epithelial cells (**Figure 4**). An Extra-Sum-of-Squares F-test demonstrated that the regression lines computed for HUMEC, BT474, and MCF7 cells were statistically different (p<0.0001). The decrease in cell number did not correspond to the rate of cell proliferation for each of the cells studied, with the HUMEC being the slowest dividing cells, the BT474 HER2+ tumor cells being in the middle, and the MCF7 tumor cells proliferating the fastest. This outcome is unexpected given the role HER2 plays in mediating cell proliferation and the amplification of HER2 expression in the BT474 cells [32]. The inhibitor concentration versus normalized dose response curves predicts an IC_50_ of 4.998-mM for the HUMEC cells, 2.835-mM for the MCF7 tumor cells, and 1.06-mM for the BT474 tumor cells.

**Figure 4:**
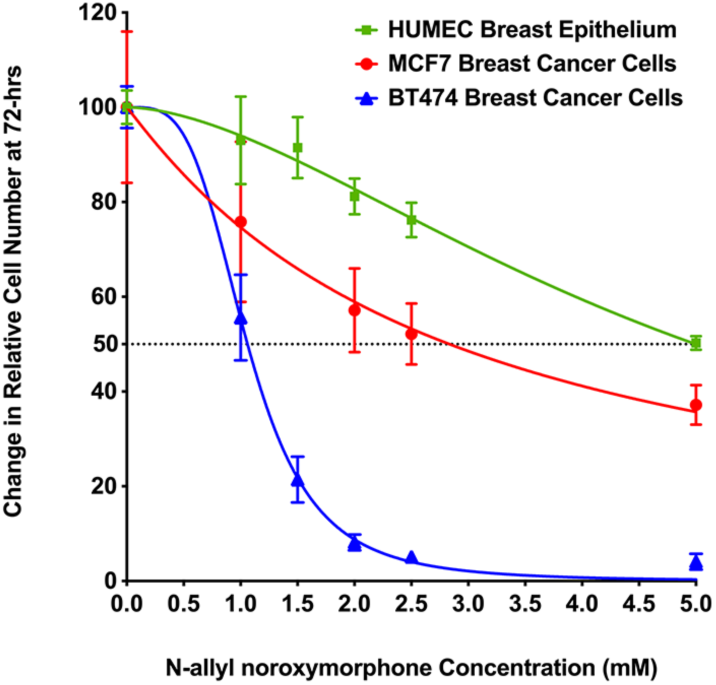
Inhibitor versus the normalized dose response cures. Non-linear regression curves were computed for the HUMEC breast epithelium (green line), MCF7 HR-positive/HER2-negative breast cancer cells (red line), and the BT474 HR-positive/HER2-positive (blue line). The regression lines were shown to be significantly different via an Extra-Sum-of-Squares F-test (p<0.0001), which means the data are best represented as separate lines. The R^2^-value was 0.9188 for the HUMEC cells, 0.7596 for the MCF7 tumor cells, and 0.9847 for the BT474 tumor cells. The IC_50_ (dashed line) was 4.998-mM for the HUMEC cells, 2.835-mM for the MCF7 tumor cells, and 1.06-mM for the BT474 tumor cells.

Extracellular LDH measured in the MCF7 cells at 72-hours after the addition of N-allyl noroxymorphone did not result in significant differences from the control cultures. These LDH values suggest that the cytotoxic mechanism does not become a significant driver of cell death until the dose is >6-mM (**Figure 5A**). Only the 8-mM and 10-mM N-allyl noroxymorphone concentrations demonstrated significant LDH expression when compared to controls (* = p<0.036 and **** = p<0.0001). This suggests that the decrease in MCF7 tumor cell number is not directly coupled to the mechanism that mediates cell death. In contrast, 24-hours after treating the BT474 cultures with N-allyl noroxymorphone, the LDH values increased steeply which corresponded to the steep decrease cell number. This demonstrates that the loss in BT474 tumor cells number is more tightly coupled with the mechanism that drives cell death (**Figure 5B**). A one-way ANOVA demonstrated that all the tested N-allyl noroxymorphone concentrations in the BT474 cultures produced a significant increase in LDH expression relative to the control cultures (* = p<0.0334, ** = p<0.0097, and **** = p<0.0001).

**Figure 5:**
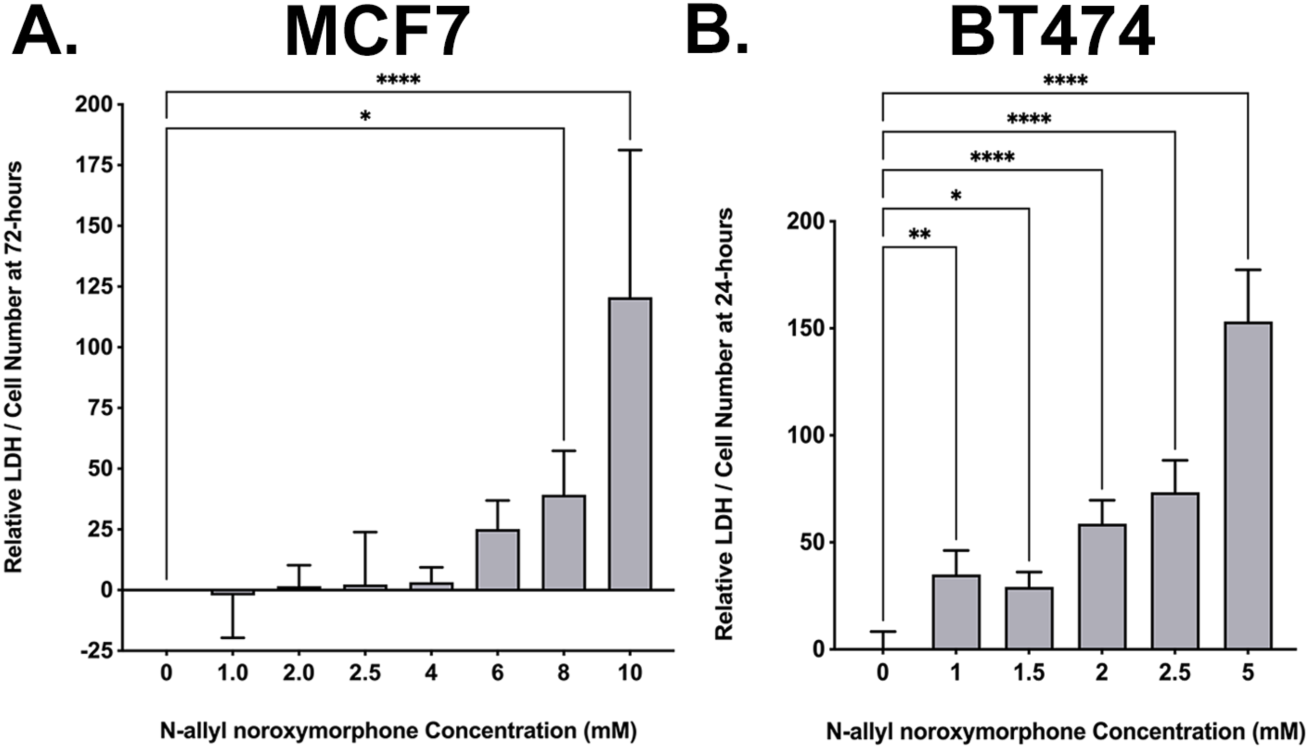
LDH expression is a surrogate for cell death. (A) MCF7 tumor cells were assessed at 72-hours following the administration of N-allyl noroxymorphone (0-, 1-, 2-, 2.5-, 4-, 6-, 8-, and 10-mM). A one-way ANOVA demonstrated that all the tested N-allyl noroxymorphone concentrations tested in MCF7 cultures produced a significant increase in LDH expression at 8- and 10-mM doses relative to controls (* = p<0.0334, **** = p<0.0001). (**B**) BT474 tumor cells were assessed at 24-hours following the administration of N-allyl noroxymorphone (0-, 1-, 1.5-, 2.5-, and 5-mM). A one-way ANOVA demonstrated that all the tested N-allyl noroxymorphone concentrations tested in BT474 cultures produced a significant increase in LDH expression relative to the control cultures (* = p<0.0334, ** = p<0.0097, and **** = p<0.0001).

The spheroid assay was used to assess the effects that the N-allyl noroxymorphone on tumor cell growth and survival. After 336-hours, crystal violet staining in MCF7 cultures decreased at the highest doses tested, 5- and 10-mM (**Figure 6A**). The area of the spheroid micro-mass decreased 41.9% for the 5-mM and 68.1% for the 10-mM N-allyl noroxymorphone concentrations, which were significantly different (* = p<0.014 and **** = p<0.0001) versus the control cultures. Following treatment with N-allyl noroxymorphone at 120-hours, a significant decrease in the spheroid micro-mass area was observed in BT474 cultures (**Figure 6B**). Treatment with the following N-allyl noroxymorphone concentrations reduced micro-mass size relative to controls cultures: 1.5-mM of N-allyl noroxymorphone reduced the micro-mass size 14.7%, 2-mM of N-allyl noroxymorphone concentration decreased the micro-mass size 23.8%, 2.5-mM of N-allyl noroxymorphone concentration decreased the decreased micro-mass size 30.6%, and 5-mM of N-allyl noroxymorphone concentration decreased the micro-mass size 84% (* = p<0.017, *** = p<0.0002, and **** = p<0.0001). The MCF7 micro-masses at 336-hours were smaller than their corresponding control cultures at that same time-point. Micro-masses in the BT474 cultures were both smaller than their corresponding control cultures, but also smaller in size than they were at the start of the experiment on day 0 (i.e., the micro-masses shrank). The spheroid assay is a useful tool to assess if animal studies are likely to be successful by testing the effects of a therapeutic agent on long-term tumor growth and is a surrogate for post-therapeutic survival. The results of the spheroid assays, in both the MCF7 and BT474 tumor cells, suggested that the effects of MBC-005 treatment would not be short-term and would negatively impact tumor survival when tested *in vivo*.

**Figure 6:**
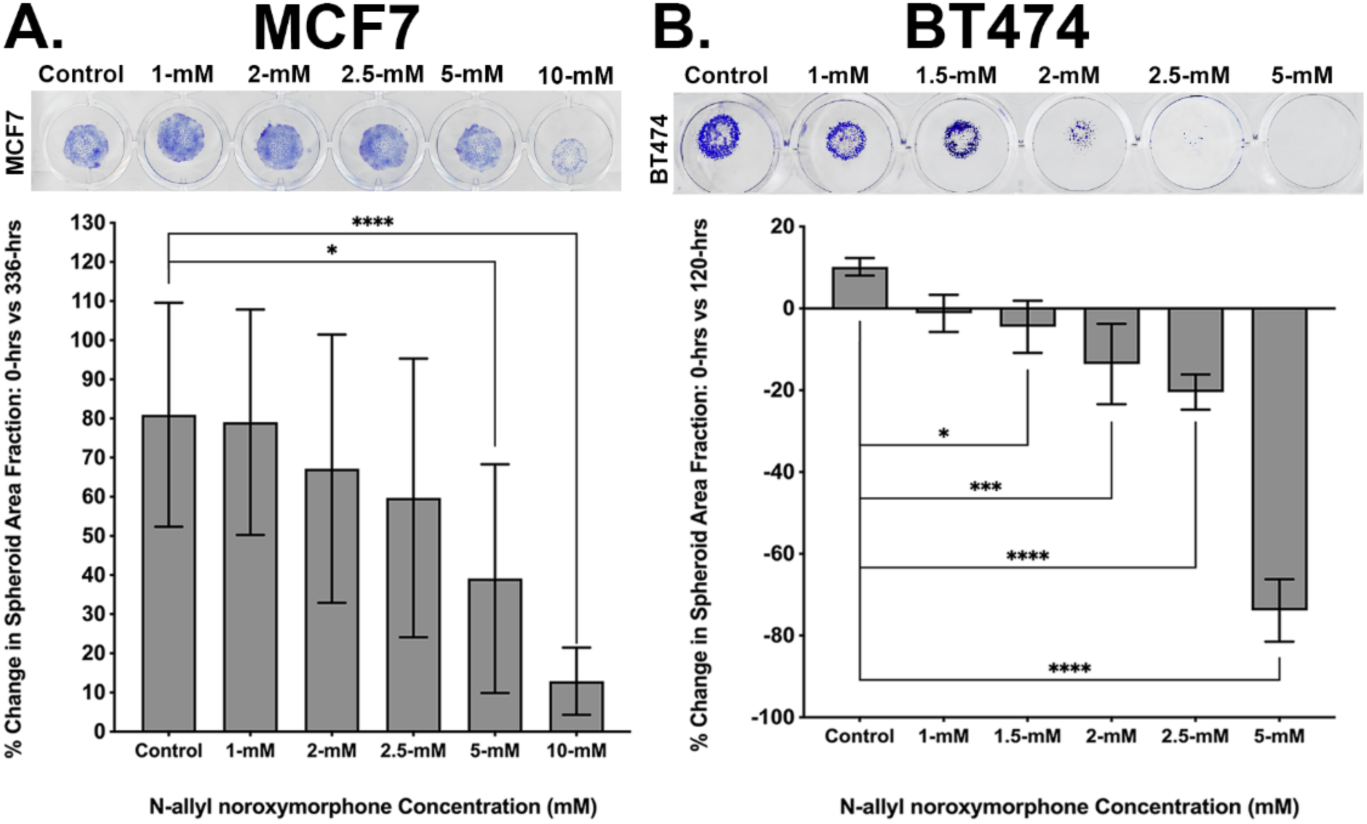
The spheroid assay is used to assess long-term effects of therapy on tumor cell by culturing the tumor cells in a micro-mass, which is a 3D tumor mass used as a surrogate for in vivo tumor cell growth dynamics. (**A**) MCF7 tumor cell micro-mass cultures (purple staining) treated with 0-, 1-, 2-, 2.5-, 5-, and 10-mM N-allyl noroxymorphone doses, after 336-hours, produced a significant decrease of 41.9% and 68.1% in micro-mass areas for the 5- and 10-mM N-allyl noroxymorphone concentrations, respectively (* = p<0.014 and **** = p<0.0001). **(B)** BT474 tumor cell micro-mass cultures (purple staining) treated with 0-, 1-, 1.5-, 2-, 2.5-, and 5-mM N-allyl noroxymorphone concentrations, after 120-hours, resulted in a significant decrease of 14.7%, 23.8%, 30.6%, and 84% for the 1.5-, 2-, 2.5, and 5-mM N-allyl noroxymorphone concentrations, respectively (* = p<0.017, *** = p<0.0002, and **** = p<0.0001).

The hydrogel component of MBC-005 was also tested for its effects on cell proliferation and cell death using the MTT and LDH assays, corrected for cell number to allow for the comparison between different cell-lines. The MC3T3-E1 (MC3) cell-line was used as non-tumor cell control. Following treatment with the hydrogel component of MBC-005, no significant change in MC3 cell number at 24- or 72-hours (**Figure 7A**). Relative to the MTT values for the MC3 cells at 24-hours, in MCF7 cultures there was a significant 6.41-fold decrease in cell number at 24-hours (p<0.0003) and a 10.41-fold decrease in cell number at 72-hours (p<0.0001). Similarly, when compared to the MTT values for the MC3 cell at 24-hours, the MTT values for the BT474 cells were significantly decreased 2.89-fold (p<0.0019) at 24-hours and 4-fold at 72-hours (p<0.0005). In addition, after treatment with the hydrogel component of MBC-005, no significant change in MC3 cell death at 24- or 72-hours (**Figure 7B**). However, relative to the LDH values for the MC3 cell at 24-hours, in MCF7 cultures there was a significant 3.15-fold increase in LDH (p<0.0001) at 24-hours and a 11-fold increase in LDH (p<0.0044) at 72-hours. For BT474 cultures there was a significant 2.8-fold increase in LDH (p<0.0001) at 24-hours and 2.62-fold at 72-hours.

**Figure 7:**
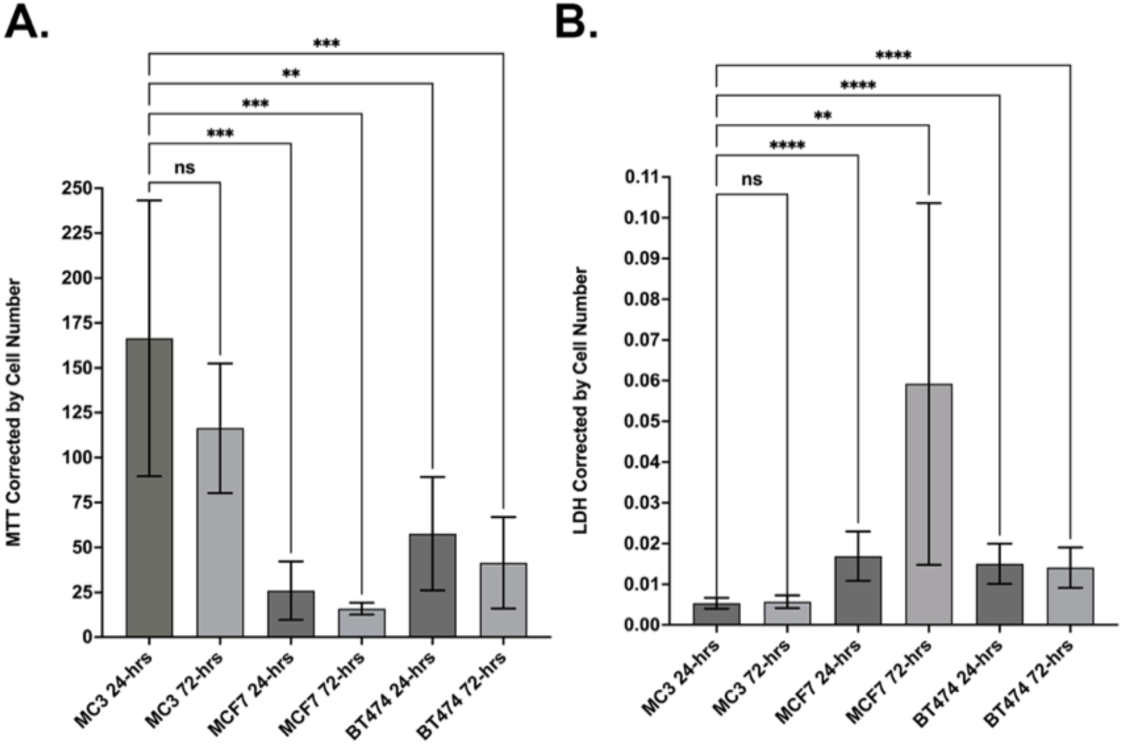
The MTT assay and the LDH assay were used to assess the effects of the hydrogel component of MBC-005 on tumor cell number and cell death. MC3T3-E1 cells (MC3) were used as a non-tumor cell control. (**A**) There was no statistically significant difference (NS) in MTT values between the MC3 cell cultures at 24- and 72-hours. Relative to the MTT values for the MC3 cells at 24-hours, in MCF7 cultures there was a significant decrease in cell number at 24-hours (*** = p<0.0003) and 72-hours (*** = p<0.0001). MTT values for the BT474 cells were significantly decreased at 24-hours (** = p<0.0019) and 72-hours (*** = p<0.0005). (**B**) There was no significant difference (NS) between LDH values for MC3 cultures at 24- and 72-hours. Relative to the LDH values for the MC3 cells at 24-hours, in MCF7 cultures there was a significant increase in LDH at 24-hours (*** = p<0.0001) and 72-hours (** = p<0.0044). In BT474 cultures LDH values were significantly increased at 24-hours (*** = p<0.0001) and 72-hours (p<0.0001).

A p21 ELISA assay was used to assess p21 protein expression levels in MCF7 and BT474 cells (**Figure 8**) following treatment with MBC-005. The p21 protein expression was normalized for total protein was interpolated using the included standards. Twenty-four hours after the addition of 1-mM MBC-005 to MCF7 cultures, p21 expression significantly increased 3.75-fold (p<0.0082) relative to control cultures (**Figure 8A**). Correspondingly, 24-hours after the addition of 1-mM MBC-005 to BT474 cultures, p21 expression significantly increased 5.78-fold (p<0.0435) relative to control cultures (**Figure 8B**).

**Figure 8:**
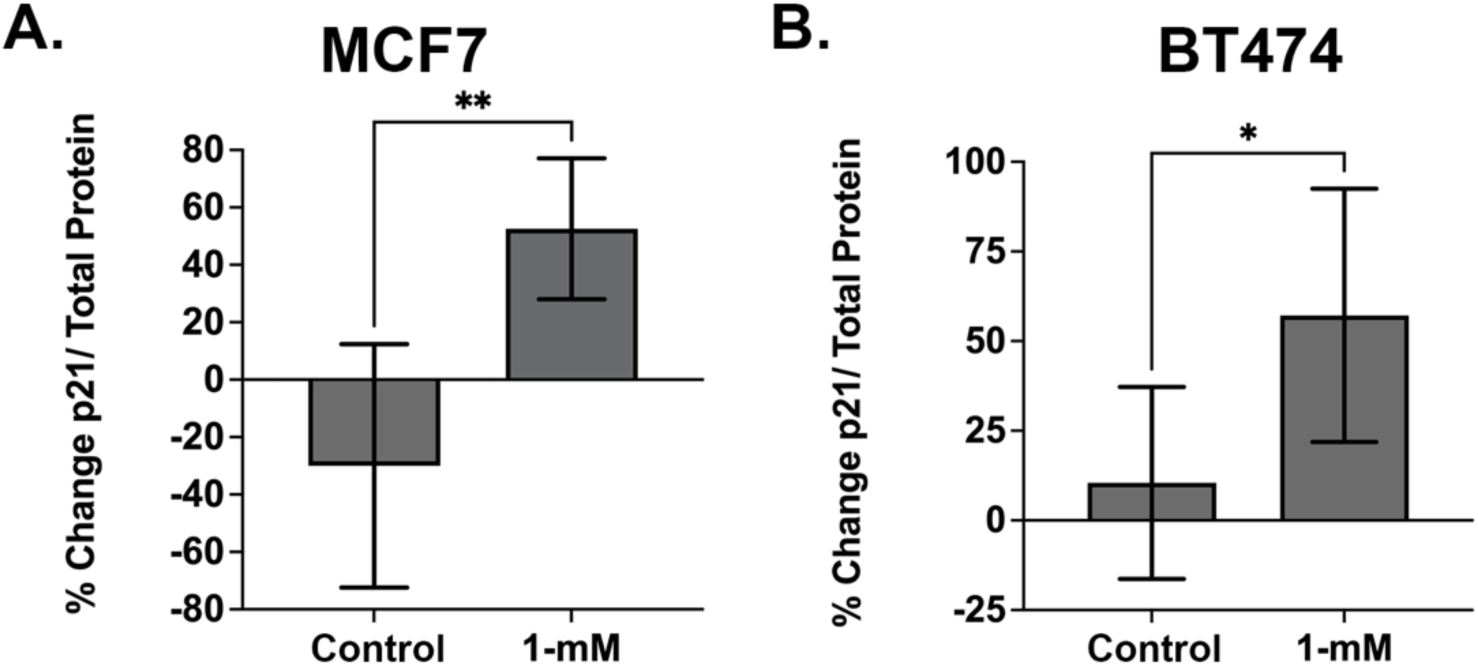
The p21 ELISA assay was used to assess the effects of MBC-005 on tumor cell expression of p21. (**A**) MCF7 tumor cells were treated with 1-mM MBC-005 and then assessed for p21 expression 24-hours later. p21 expression increased significantly 3.75-fold (** = p<0.0082) relative to control cultures. (**B**) BT474 tumor cells were also treated with 1-mM MBC-005 and then assessed for p21 expression 24-hours later. p21 expression increased significantly 5.78-fold (* = p<0.0435) relative to control cultures.

MBC-005 was also tested versus doxorubicin in MCF7 tumor cells (**Figure 9**). Relative to control cultures, 24-hours after adding 1-mM and 10-mM of MBC-005, cell numbers decreased 24.2% and 65.1%, respectively (p<0.0001). Additionally, the 10-mM MBC-005 dose was also shown to be significantly decreased 53.9% than the 1-mM MBC-005 dose (p<0.0001). Cell number increased after the addition of 150-nM doxorubicin 4.5%, although this change in cell number was not statistically significant. In contrast, the 150-μM concentration of doxorubicin resulted in a significant 78.8% decrease in cell number (p<0.0001). Interestingly, the 10-mM MBC-005 dose and the 150-μM doxorubicin dose were not significantly different.

**Figure 9:**
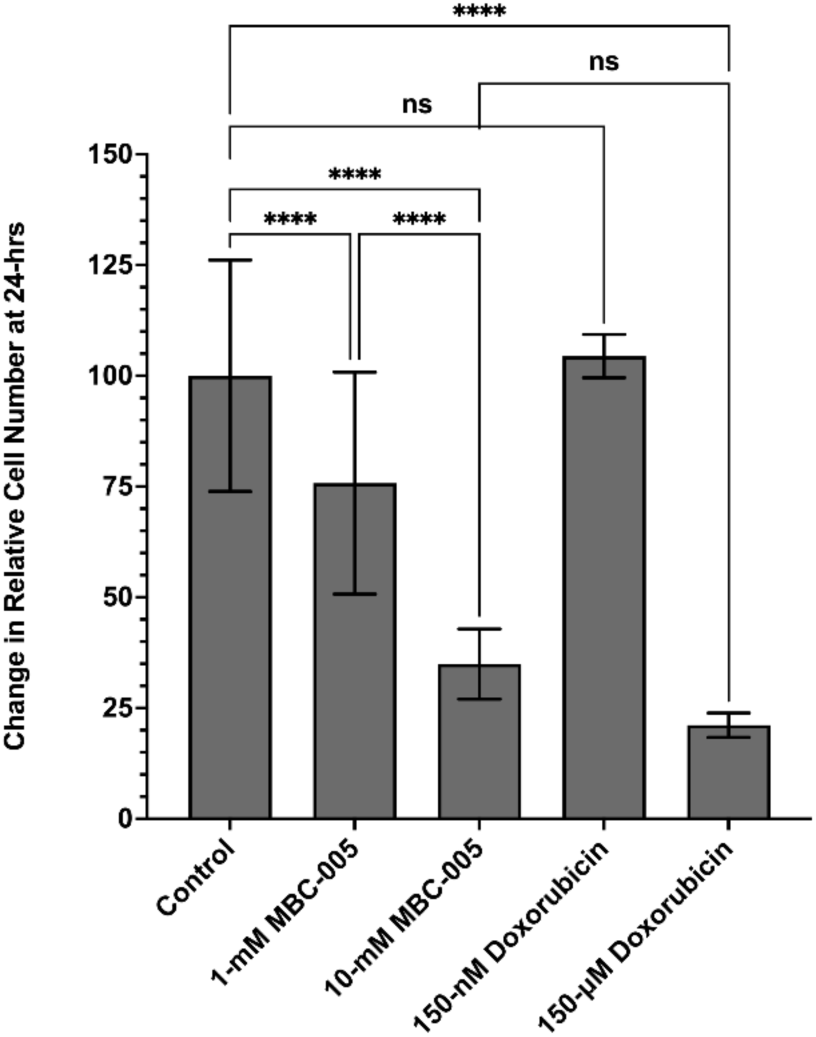
MCF7 cultures were treated with either MBC-005 or doxorubicin. Treatment with 1-mM MBC-005 or 10-mM MBC-005 produced a significant decrease in cell number of 24.2% and 65.1%, respectively, 24-hours after treatment (**** = p<0.0001). A further significant decreased of 53.9% was observed between the 1-mM MBC-005 dose and the 10-mM MBC-005 dose (**** = p<0.0001). Treatment with 150-nM of doxorubicin did not produce a significant change in cell number relative to control cultures (NS). Cultures treated with 150-μM of doxorubicin were seen to significantly decrease 78.8% relative to controls (**** = p<0.0001). Comparison between the 10-mM MBC-005 dose and the 150-μM dose of doxorubicin were also found to be not significant (NS).

### MBC-005 Increased Survival in a Mouse Xenograft Liver Metastases Model

The 4T1 breast cancer cell line, tagged with luc2 luciferase (4T1-luc2), was implanted directly into the liver of BALB/c mice (N=12 per treatment group). Seven days after tumor inoculation, mice were treated with various concentrations of MBC-005 (30-, 60-, 120-, 180-, 240-, or 480-μg) in combination with 5-mg/kg doxorubicin every 3-days. Mice in the Control MBC-005 group were treated with MBC-005 that lacked N-allyl noroxymorphone in combination with 5-mg/kg doxorubicin every 3-days. Mice in the No Treatment Control (NTC) group did not receive any therapy. Animals were then observed over 23-days using bioluminescent imaging on day 0, 8, and 15 after treatment with MBC-005. A Cox proportional hazards model was constructed using the following: 1) treatment versus no treatment, 2) the presence of lung metastases, 3) the calculated area under the curve (AUC) for the whole-body bioluminescence as a surrogate for tumor burden, and 4) the change in body weight as a surrogate for overall animal health. The presence of lung metastases and the change in body weight had the greatest impact on the model (**Table I**). When compared to the NTC group, the 60-μg MBC-005 dose group had the most significant survival benefit, with an HR = 0.4471. Further, interpolated cumulative survival (**Table II**) was also greatest in the mice treated with 60-μg MBC-005, both when they did not have lung metastases (52.67%) and then they did have lung metastases (3.07%).

**Table I:**
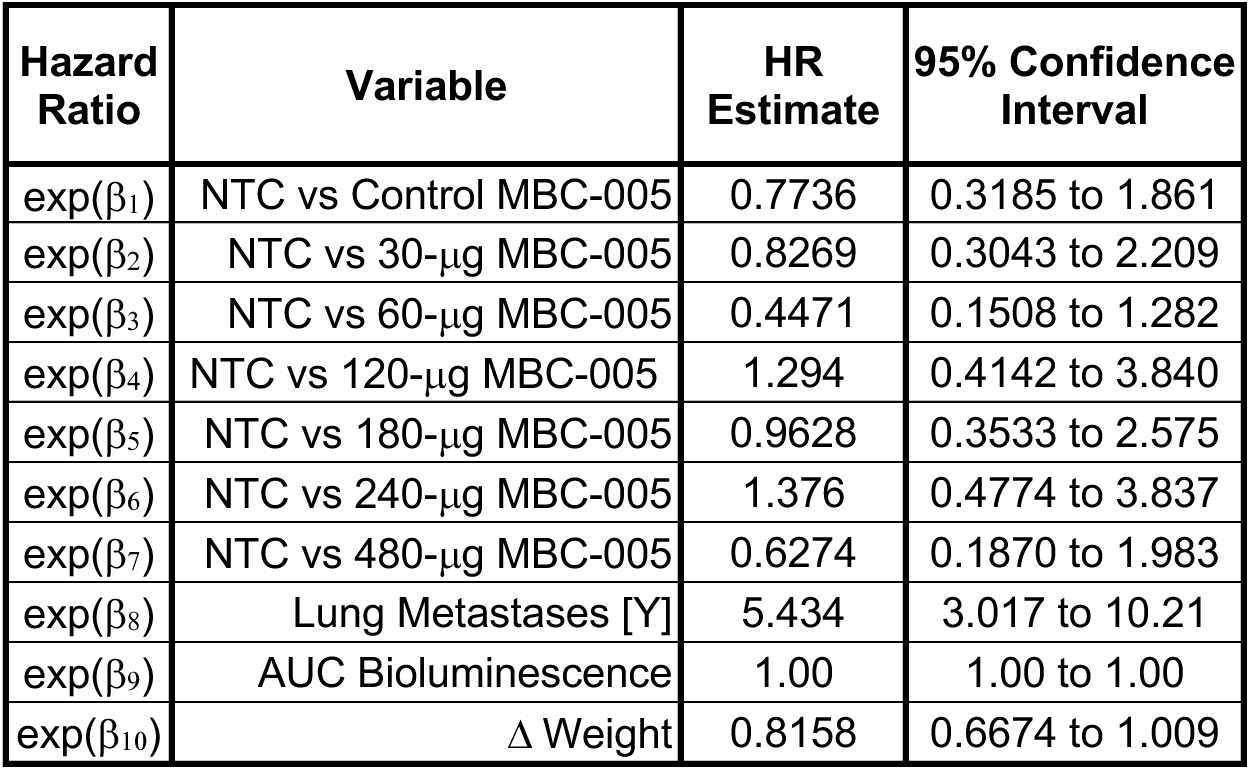
Cox Proportional Hazard Ratios.

| Hazard Ratio | Variable | HR Estimate | 95% Confidence Interval |
| --- | --- | --- | --- |
| $\exp(\beta_1)$ | NTC vs Control MBC-005 | 0.7736 | 0.3185 to 1.861 |
| $\exp(\beta_2)$ | NTC vs 30-μg MBC-005 | 0.8269 | 0.3043 to 2.209 |
| $\exp(\beta_3)$ | NTC vs 60-μg MBC-005 | 0.4471 | 0.1508 to 1.282 |
| $\exp(\beta_4)$ | NTC vs 120-μg MBC-005 | 1.294 | 0.4142 to 3.840 |
| $\exp(\beta_5)$ | NTC vs 180-μg MBC-005 | 0.9628 | 0.3533 to 2.575 |
| $\exp(\beta_6)$ | NTC vs 240-μg MBC-005 | 1.376 | 0.4774 to 3.837 |
| $\exp(\beta_7)$ | NTC vs 480-μg MBC-005 | 0.6274 | 0.1870 to 1.983 |
| $\exp(\beta_8)$ | Lung Metastases [Y] | 5.434 | 3.017 to 10.21 |
| $\exp(\beta_9)$ | AUC Bioluminescence | 1.00 | 1.00 to 1.00 |
| $\exp(\beta_{10})$ | Δ Weight | 0.8158 | 0.6674 to 1.009 |

**Table II:** Interpolated survival in mice with or without lung metastases.

| Treatment Group | Survival with No Lung Metastases |  | Survival with Lung Metastases |  |
| --- | --- | --- | --- | --- |
|  | Interpolated Cumulative Survival | Interpolated HR | Interpolated Cumulative Survival | Interpolated HR |
| NTC | 23.84% | 0.582 | 0.04% | 3.160 |
| Control MBC-005 | 32.98% | 0.450 | 0.20% | 2.445 |
| 30-μg MBC-005 | 30.55% | 0.481 | 0.16% | 2.613 |
| 60-μg MBC-005 | 52.67% | 0.260 | 3.07% | 1.413 |
| 120-μg MBC-005 | 15.63% | 0.753 | 0.004% | 4.090 |
| 180-μg MBC-005 | 25.14% | 0.560 | 0.06% | 3.043 |
| 240-μg MBC-005 | 13.90% | 0.800 | 0.002% | 4.349 |
| 480-μg MBC-005 | 40.67% | 0.365 | 0.08% | 1.983 |

Overall survival in the NTC group mice was 8.5% at day 23 (**Figure 10**). Control MBC-005 group survival was 14.8%, which is 74% greater than NTC group survival, which may be related to doxorubicin treatment. The 60-μg MBC-005 treatment group was 3.9-fold greater than the NTC group and 2.24-fold greater than the Control MBC-005 treatment group. These results suggest that the 60-μg MBC-005 dose group has a significant survival benefit over 5-mg/kg doxorubicin.

**Figure 10:**
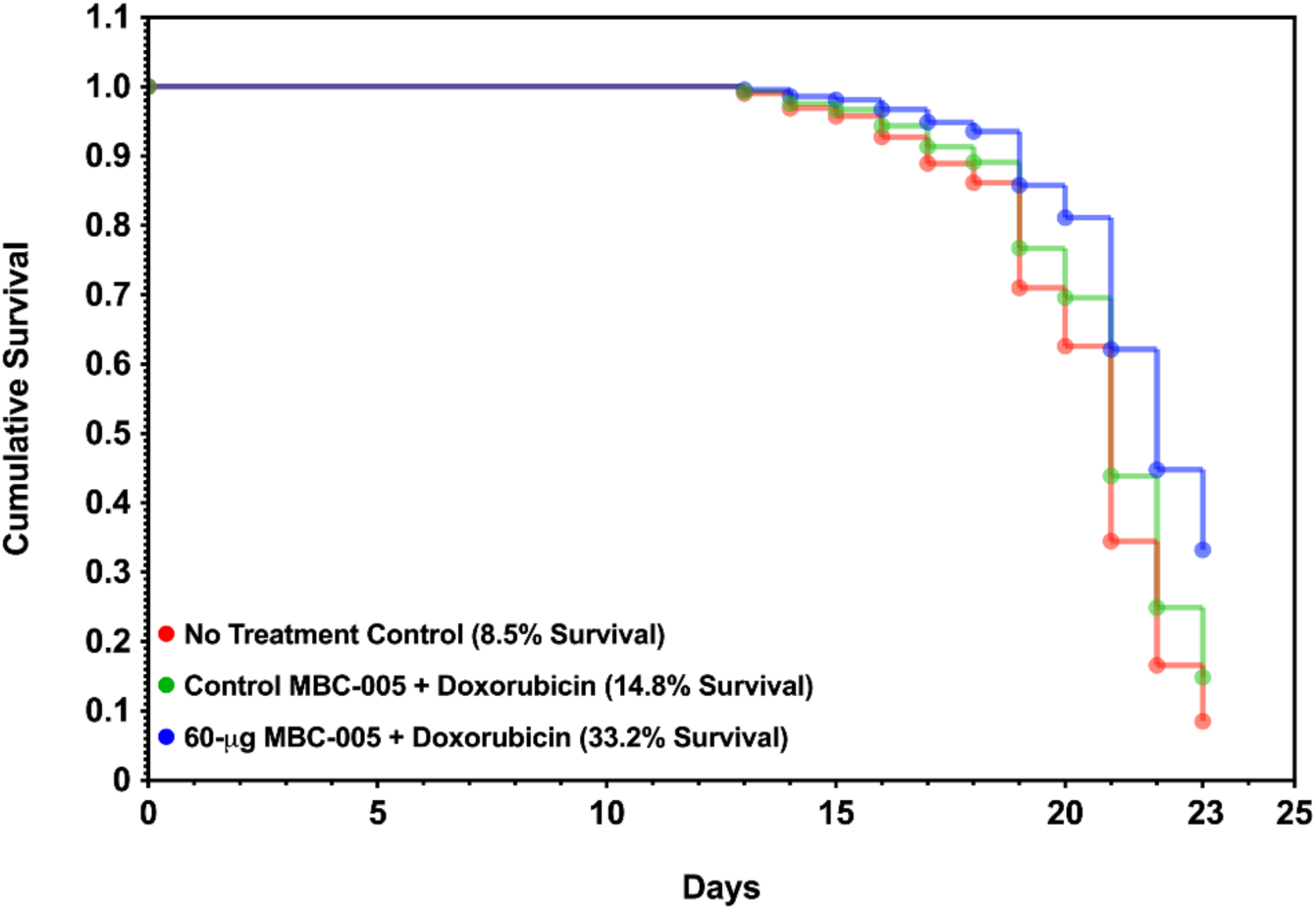
Survival in Mice with Liver Metastases after MBC-005 Treatment. Survival in BALB/c mice inoculated with 4T1-luc2 breast cancer cells directly in the liver were assessed using a Cox proportional hazard model. At day 23, 8.5% of the mice in the No Treatment Control group were observed to have survived (red). The survival in the Control MBC-005 treatment group mice was 14.8% and the survival in the 60-μg MBC-005 group was 33.2%.

The 4T1-luc2 breast cancer tumor cells can be tracked using bioluminescence, which corresponds to tumor burden. A Malthusian exponential growth model was developed using non-linear least-squares regression to estimate the doubling time for the tumor bioluminescence [33]. Regression lines were then assessed for statistical differences between groups using an Extra-Sum-of-Squares F-test to determine if the groups were statistically different or, alternatively, could be modeled using a single regression line (α=0.05). Using this method to assess the whole-body bioluminescent signal all the groups were found to be significantly different from one another (p<0.0001) (**Figure 11A**). The same analytical approach was used to assess the bioluminescent signal from the liver, and all the groups were found to be significantly different from one another (p<0.0001) (**Figure 11B**). In both the whole-body and in the liver the bioluminescent signal in the 60-μg MBC-005 treatment group was the lowest. The Malthusian exponential growth model also allows for the determination of the doubling-time, which corresponds to time-dependent tumor growth. The whole-body bioluminescent signal doubling-time for the NTC group was 5.7-days (**Table III**). Relative to the NTC group, all the MBC-005 treatment groups had a significantly increased doubling time. In particular, the 60-μg MBC-005 dose group had the longest doubling-time of 27-days, which corresponds to the slowest tumor growth of the treatment groups.

**Figure 11:**
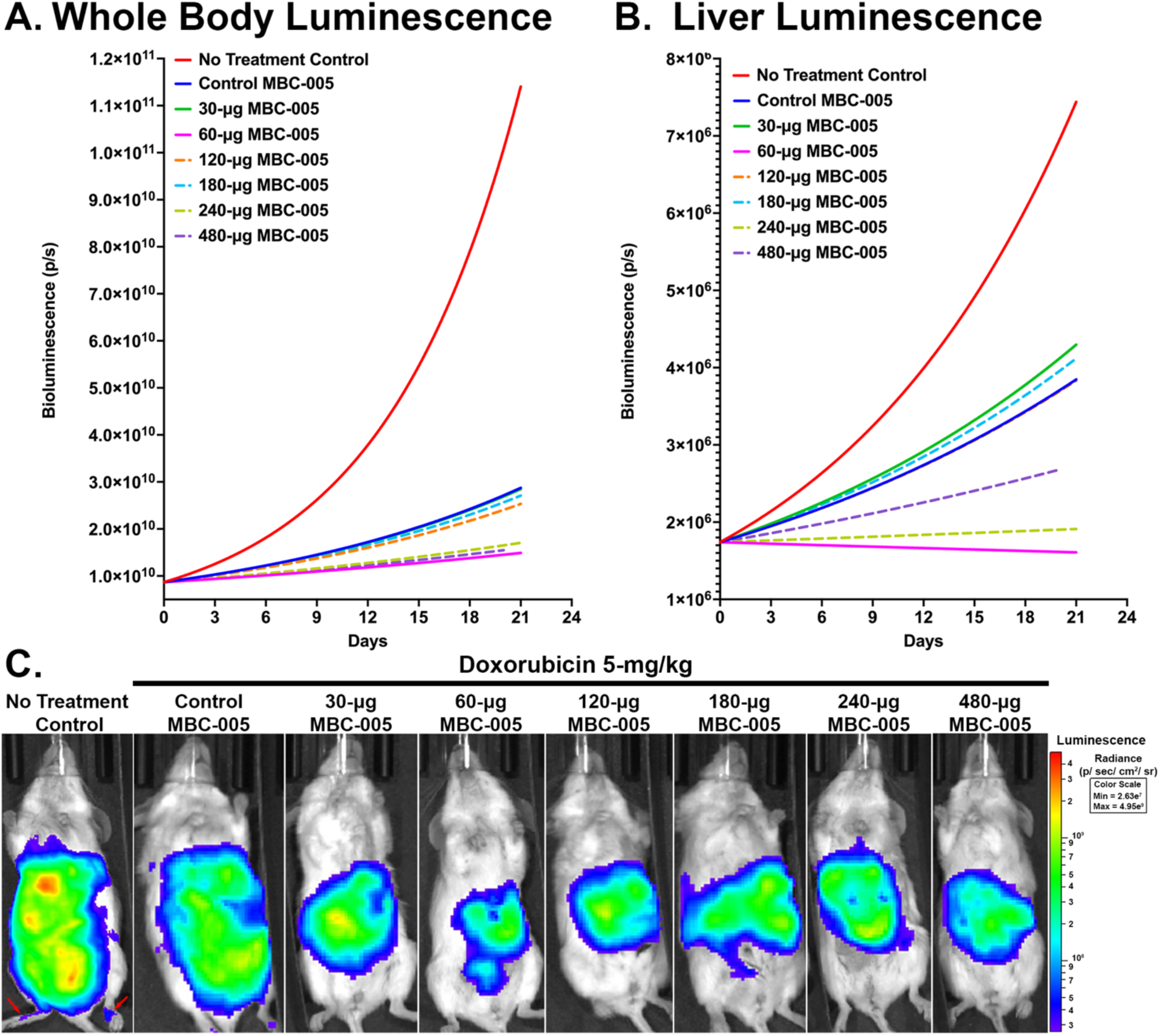
Mice in the NTC and MBC-005 treatment groups were evaluated for bioluminescent signaling at 6-, 13-, and 21-days after tumor inoculation. (**A**) Malthusian non-linear regression analysis demonstrated that the whole-body bioluminescent signal was significantly different between the different treatment groups (p<0.0001). (**B**) A Malthusian non-linear regression analysis demonstrated that the liver bioluminescent signal was significantly different between the treatment groups (p<0.0001). (**C**) Bioluminescent imaging showed the greatest decrease in signal was in the 60-μg MBC-005 groups; however, 30-, 120-, 180-, 240-, and 480-μg MBC-005 groups qualitative had less bioluminescent signal. In addition, the NTC group animal had bioluminescent signal in its hind-limb (red arrow).

**Table III:** Bioluminescent Doubling Time.

| Treatment Groups | Doubling Time (days) |
| --- | --- |
| NTC | 5.7 |
| Control MBC-005 | 12.2 |
| 30-μg MBC-005 | 12.3 |
| 60-μg MBC-005 | 27.0 |
| 120-μg MBC-005 | 13.6 |
| 180-μg MBC-005 | 12.8 |
| 240-μg MBC-005 | 21.6 |
| 480-μg MBC-005 | 24.0 |

The bioluminescence was lowest in the mice in the 60-μg MBC-005 treatment group (**Figure 11C**). In addition, the 30-, 120-, 180-, 240-, and 480-μg MBC-005 groups also had less bioluminescent signal than the NTC group and the Control MBC-005 groups, the latter of which had widely disseminated tumor throughout the abdominal and thoracic cavities. Further, at 14-days 58.3% (7/12) of the mice in the NTC group had metastases in their hind-limbs and at 21-days the one remaining mouse in the NTC group also had metastases in its hind-limb (**Figure 11C**). None of the animals treated with MBC-005 had hind-limb metastases. Post-hoc analysis using 2-way ANOVA was used to evaluate differences between dose treatment groups.

An analysis of liver weights corrected for whole body weight (%liver weight) was assessed as a surrogate for liver tumor burden (**Figure 12**). Mean values of the %liver weight falling within the highlighted bar are not significantly different than the reference control values. Administration of 60-µg MBC-005 resulted in the lowest %liver weight. Clinical chemistry and hematology were assessed in mice at day 23, the conclusion of the study. Alkaline phosphatase (ALP) is a liver enzyme that can relate to liver stress. ALP expression was in the normal range for the 60- and 120-µg MBC-005 treatment groups. Alanine aminotransferase and aspartate aminotransferase are liver enzymes that can also indicate liver stress. However, no clear trend was observed for the different treatment groups. Likewise, the assessment of hematological values also did not show any treatment related pattern.

**Figure 12:**
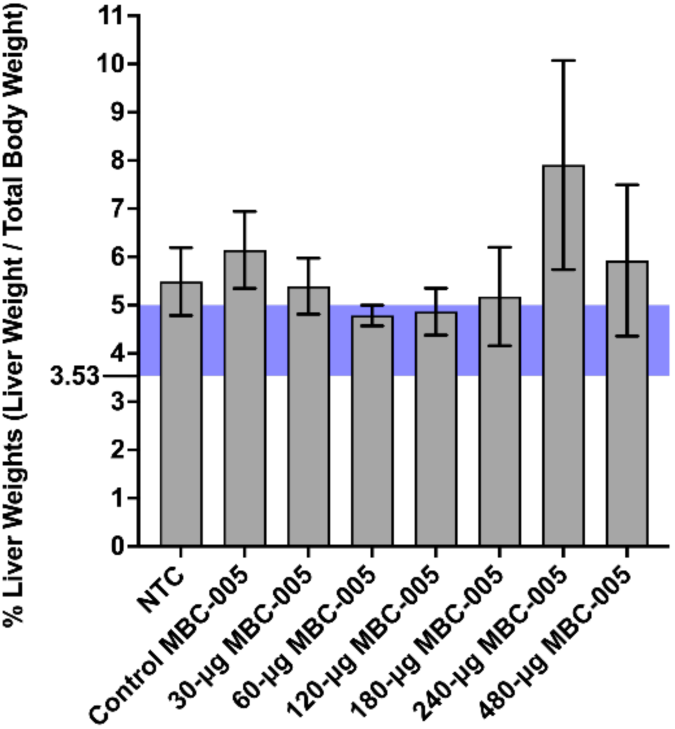
Liver weights were normalized by the mouse body weight to derive the %liver weight. Reference %liver weight was derived from published reference liver and total body weights and ranged from 3.53-mg to 5-mg. Only the 60- and 120-μg MBC-005 treatment groups were within the reference range, with means of 4.785-mg (SEM ±0.2144) and 4.863-mg (SEM ±0.4863), respectively. Only the 60-μg MBC-005 treatment group has an SEM that is below 5-mg.

Liver samples were assessed for tumor volumes and necrosis volumes using histology (**Figure 13**). Histomorphometric analysis demonstrated that administration of 60-µg MBC-005 resulted in a significant 4.2-fold decrease (p<0.0038) in the liver volume fraction (V_f_) relative to the NTC group, which is a morphometric feature that is directly related to the tumor volume in the liver (**Figure 14A**). Administration of 120-µg MBC-005 also resulted in a significant (p<0.0155) decrease of 2.3-fold in tumor burden in the liver relative to the NTC group. However, none of the other concentrations of MBC-005 tested resulted in a significant decrease in tumor volume. Necrosis was also histomorphometrically assessed in liver samples. No significant change in necrosis was observed (**Figure 14B**). While not significant, treatment with 60-µg MBC-005 resulted in a 3-fold decrease in necrosis while administration with 120-µg MBC-005 resulted in a 2.25-fold decrease in necrosis. Fibrosis was also significantly greater in the MBC-005 treated animals when compared to the animals in the NTC group. This fibrotic tissue was often observed to have small clusters of hepatocytes. This study wasn’t designed to assess the residual the effects of MBC-005. However, in liver samples in which residual MBC-005 remained, there was no evidence of adverse tissue reactivity. In the 60-µg MBC-005, 120-µg MBC-005, and 480-µg MBC-005 group samples in which residual MBC-005 can be observed (red arrows), it appears adjacent to healthy liver tissue.

**Figure 13:**
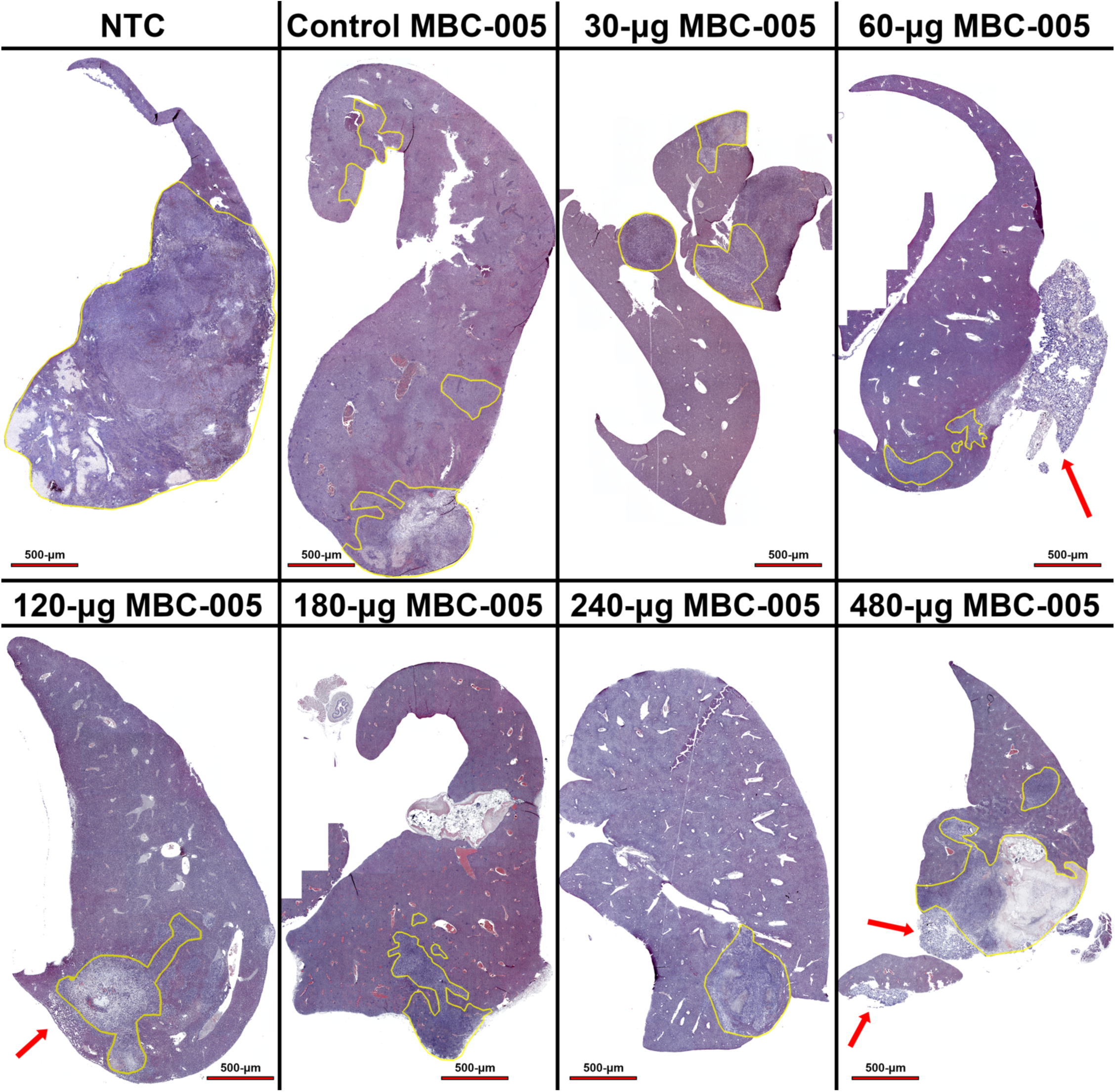
Liver samples were stained with hematoxylin and eosin and imaged at 4x. Tumor is highlighted in the yellow. outline Residual MBC-005 (red arrows) was adjacent to healthy liver tissue as well as tumor.

**Figure 14:**
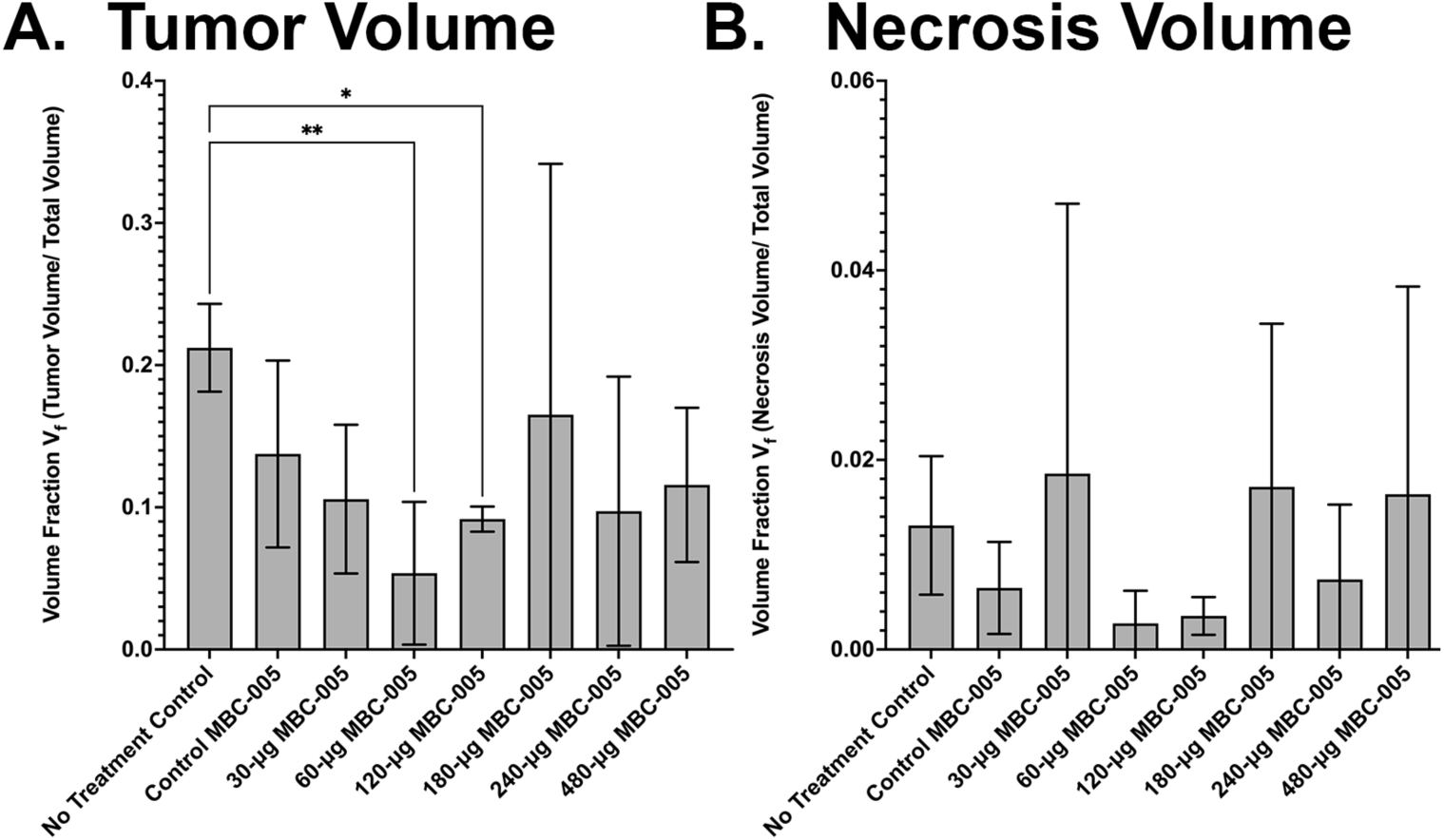
Histomorphometric analysis of the tumor volume fraction (V_f_) in the liver and necrosis volume fraction. (**A**) Treatment with 60-mg MBC-005 resulted in a 4.2-fold decrease in tumor volume fraction (** = p<0.0038). Treatment with 120-mg MBC-005 resulted in a 2.3-fold decrease in tumor volume fractions (* = p<0.0155). (**B**) Necrosis volume fraction was not significantly different between groups; however, there was a 3-fold decrease in necrosis for the 60-mg MBC-005 and a 2.25-fold decrease for the 120-mg MBC-005 treatment groups

## Discussion

There are many limitations associated with the administration of small molecules for use as anti-cancer drug therapy, which include hydrophobicity, solubility, enzymatic degradation, off-target toxicity, and renal clearance [1–4]. In previous work, we identified that N-allyl noroxymorphone had significant therapeutic benefit to inhibit cell proliferation via a p21-dependent mechanism [7]. N-allyl noroxymorphone has two properties that limit its potential therapeutic utility: it is hydrophobic at pH >7 and it has a half-life of 60-minutes due to enzymatic degradation [5, 6]. These properties make it difficult to achieve sustained serum concentrations of N-allyl noroxymorphone that are high enough to have a therapeutic effect. This is particularly true for tissues with low rates of perfusion, such as cartilage or bone. In our previous work, we overcame these limitations in bone by employing a direct injection methodology to deliver the N-allyl noroxymorphone [7, 34]. Clinical studies to treat bone metastases in the spine via a local administration of N-allyl noroxymorphone are ongoing (NCT05280067).

The N-allyl noroxymorphone formulation delivered directly to malignancies in bone is novel and designed specifically to treat breast cancer metastases in bone [34]. Breast cancer metastases in bone are destructive, resulting in fracture and produce debilitating pain. Recent experience with a single patient treated via FDA’s Expanded Access program (i.e., also known as compassionate use) resulted in an encouraging clinical outcome, in which percutaneous local administration of the N-allyl noroxymorphone formulation was injected directly into the metastases located in the vertebral bone. This approach was able to successfully treat the three separate breast cancer metastases in this patient’s spine, resulting in resolution of the tumor lesions and significantly decreased pain. This patient has remained free of tumor in the treated vertebral bodies 3-years following the local, percutaneous administration of the N-allyl noroxymorphone. This result provided the impetus to examine if N-allyl noroxymorphone could be used to treat breast cancer metastases in other parts of the body, and particularly the liver.

Breast cancer metastases in soft tissues are common in liver, lung, and the brain; however, liver metastases are more amenable to the local administration of therapy due to their anatomic location and the ability to image the metastases. Patients with disseminated metastatic disease have liver involvement in 50% to 62% of cases [11, 12]. Adam et al. reported a median survival of between 4- to 8-months for patients without treatment [35]. Rivera et al. reported that patients treated with a combination of chemotherapy, immunotherapy, hormone therapy had a 5-year survival <12%, with median overall survival of between 4- to 21-months [13]. In general, however, survival is highly variable and is based on many factors, which include the breast cancer receptor sub-type. For instance, Xie et al. found that patients with the hormone receptor (HR)+/HER2+ subtype had a median survival of 31-months [36]. They also identified that HR-/HER-2+ patients had a median survival of 22-months, and those with triple negative breast cancer (TNBC) had median survival of 8-months. Ji et al. reported that out of 311,573 patients from the SEER database that 25.6% of patients with disseminated metastatic disease had liver metastases with a median overall survival was 20-months [37]. They also found the median overall survival times for the following subtypes: HR+/HER2+ median overall survival of 38-months, HR-/HER2+ median overall survival of ≤31-months, HR+/HER2-median overall survival 21-months, and TNBC median overall survival 9-months.

The current SOC for breast cancer liver metastases is systemic chemotherapy followed by locoregional intervention for symptomatic metastatic lesions [38]. However, the exact therapeutic interventions vary considerably and is based on the previous treatments administered earlier in the patient’s disease course, breast cancer subtype, patient demographics (e.g., age and menstruation status), and pathologic subtype (e.g., adenocarcinoma, infiltrating ductal carcinoma, and lobular carcinoma) [39]. Endocrine therapy such as aromatase inhibitors, fulvestrant or tamoxifen are used in patients with an HR+ tumor phenotype either alone [40], or more commonly in combination with any of the following kinase inhibitors, such as the CD4/6 inhibitors (e.g., palbociclib, ribociclib, or abemaciclib), the mTOR inhibitor (e.g., everolimus), or the PI3K/AKT inhibitors (e.g., alpelisib, capivasertib) [41–44]. Additionally, the newer oral selective estrogen receptor (ER) degrader (SERM) drugs elacestrant can also be used to treat HR+ liver metastases with ESR1 (estrogen receptor) gene mutations [45] Chemotherapies used to treat liver metastases includes anthracycline drugs (e.g., doxorubicin), taxane drugs (e.g., paclitaxel), alkylating agents (e.g., cyclophosphamide), fluoropyrimidine based drugs (e.g., fluorouracil [5-FU]) or the prodrug capecitabine, antimetabolic drugs such as methotrexate, vinca alkaloid agents (e.g., vinorelbine), or platinum-based drugs (e.g., carboplatin) [46–53]. Twenty percent of patients with TNBC tumors are positive for PD-L1; and, as such, patients with liver metastases can be treated with PD-1 inhibitors (e.g., pembrolizumab) with chemotherapy [54].

For patients whose liver metastases continue progress despite systemic therapies and are symptomatic, locoregional therapies are employed. These therapies include hepatic resection, but only <10% of breast cancer liver metastases are capable of this surgical intervention [55]. The median overall survival after hepatectomy has been found to be on average to 45-months and has a 5-year survival that ranges from 21% to 78% [13]. Yoon-Flannery et al. in a meta-analysis that included 1,497 patients that were treated with hepatectomy, the weighted average overall survival was 43.2-months, the 5-years survival was 36.8%, and the local recurrence was 51.4% [56]. Adam et al. in retrospective review of 85 hepatectomies found the median overall survival to be 32-months and the 5-year survival to be 37% [35]. At 10-years Adam et al. found only 16% overall survival; however, these authors also found that there was a 69% rate of recurrence with a median time to recurrence of 10-months. Given the relatively small number of patients who are eligible for hepatectomies and the high rate of liver recurrence (>50%), locoregional interventions need to be considered.

For unresectable liver metastases, ablative therapies are administered palliatively [57]. The major limitation in assessing locoregional therapies are the lack of randomized clinical trials, the small patient pool in retrospective studies, the heterogenous nature of breast cancer as a disease, and the heterogenous application of therapeutic interventions. Radiofrequency ablation (RFA) employs medium frequency alternating current to produce heat that ablates the tumor. Bai et al. reported in a retrospective study of 69 patients a median overall survival of 26-months and a 5-year survival of 11% [58]. In contrast, Meloni et al. in a prospective study found a median overall survival of 29.9-months and a 5-year survival of 32% [59]. Rivera et al. in their meta-analysis reported a median overall survival of 38-months and a 5-year survival <33% [13]. Transcatheter arterial chemoembolization (TACE) employs chemotherapy combined with beads that occlude the hepatic artery, thus depriving the tumor of blood-flow and providing a means of directly administering chemotherapy. Vogl et al. in a prospective study of 161 patients treated with TACE that contained mitomycin C/gemcitabine in combination or mitomycin C alone found a median overall survival of 32.5-months [60]. This is considerably greater than the median overall survival observed by Rivera et al. of 19.6-months based on their meta-analysis [13]. Trans arterial radioembolization (TARE) uses yttrium-90 coated glass beads or resin to deliver β-radiation to the tumor cells. Fendler et al. in a prospective series with 81 patients found that the median overall survival was approximately 10.25-months [61]. These authors also found that when the ratio of tumor volume to liver volume was less than 25% that the medial overall survival was between 37.5- and 50-months. In a meta-analysis Liu et al. found that in 412 patients that the median overall survival was 9.8-months and that this increased to 10.5-months when the ratio of tumor volume to liver volume was less than 25% [62].

Percutaneous ethanol injection (PEI) chemo-ablative technique, in which ethanol is injected directly into the tumor mass via ultrasound guidance. PEI has been used as a first-line therapy to treat hepatocarcinoma successfully. For instance, Taniguchi et al. demonstrated in a series of 31 patients had 5-year survival rates of 49.9% [63]. Interestingly, unlike other locoregional therapies, PEI can be administered on a weekly basis. Ebara et al. in a retrospective study of 270 patients found 5-year survival rates after PEI to be 60.3% [64]. These clinical study outcomes matched the 5-year survival reported in a review by McCarley et al., which were between 40% and 60% [65]. However, one of the limitations to PEI is that there is nothing to contain the ethanol once it has been injected into the tumor, which can lead to off-target damage to adjacent healthy tissues [65–67]. This has led to its application being restricted to tumors with well-defined capsules, such as hepatocarcinoma.

It is important to recognize that localized liver therapies have not been shown to improve survival in breast cancer patients within the context of randomized clinical trials. Nevertheless, in breast cancer patients currently approved locoregional therapies are typically used for symptom management, or sometimes to treat a rapidly progressing solitary liver lesion. However, locoregional therapies to treat liver metastases from colorectal carcinoma can be curative [65, 66], which has prompted continued efforts to identify interventions that might be curative for other tumor phenotypes. Further, several randomized clinical studies have identified a treatment related benefit for patients receiving stereotactic X-radiation therapy (SABR). For instance, the SABR-COMET (NCT01446744) study was a randomized clinical trial that examined if there was a clinical benefit from treating patients (N=99) with oligometastatic disease with the palliative standard of care (SOC) or administering SABR with the palliative standard of care (SABR-SOC) [68, 69]. In this study, multiple tumor phenotypes (e.g., breast, prostate, lung, and colorectal cancers) were treated at multiple different metastatic sites (e.g., liver, bone, lung, and adrenals). The study identified a significant median overall survival increase of 56% for patients treated SABR-SOC with a median overall survival of 50-months in this arm of the study versus patients treated with the SOC with a median overall survival of 28-months. This study also identified that micro-metastases present at the time of initial treatment likely drive the addition of new metastatic lesions and contributed to death. Thirty percent (30%) of patients in this study developed new metastases and received additional SABR treatments, which the authors called ‘salvage’ SABR [69]. Likewise, the NHS England study found a significant benefit from the administration of SABR to patients (N=1422) with a variety of tumor phenotypes in different metastatic locations [70], with overall survival being found to be 79.2% at 2-years. In contrast, the CURB study (NCT03808662), which was composed of patients (N=102) with non-small cell lung cancer or breast cancer, showed no benefit for breast cancer patients treated with SABR versus those treated with SOC [71–73].

The NRG-BR002 study (NCT02364557), in contrast to the above studies, was a randomized clinical study that focused only on patients with oligometastatic disease from breast cancer [74]. In this study, patients (N=125) were randomized into a SOC arm or a SABR + SOC arm. After 3-years of follow-up, there was no significant difference between the treatment arms in progression free survival or in overall survival. However, similar to the randomized trials that tested the efficacy of SABR, the NRG-BR002 study treated oligometastatic disease in a number of different tissues (e.g., bone, liver, lung); however, an analysis of the treatment effects in the different tissues in which metastases were identified has been reported. Despite these discouraging results, it is important to note that breast cancer is highly heterogenous, with different hormone receptor status, HER2 status, and proliferative rate. A limitation of the NRG-BR002 study might be that it was underpowered to accommodate the heterogeneity of breast cancer phenotypes or the additional heterogeneity in the location of metastases. It might also represent a limitation of X-radiation therapy such as SABR, which effects tumor survival through DNA strand-breaks that induce cell death via apoptosis [75, 76]. Therefore, in a study powered to examine different breast cancer phenotypes in different metastatic locations might have produced a different result. However, this reinforces the need for targeted local therapies, such as MBC-005, which circumvents some of the limitations of X-radiation therapy by inducing cell cycle arrest independent of DNA damage and apoptosis.

MBC-005 was designed to deliver our novel formulation of N-allyl noroxymorphone to breast cancer liver metastases via a percutaneous image-guided injection. Like other locoregional ablative therapies, such as PEI and RFA, MBC-005 was designed to be administered repeatedly; however, unlike PEI or RFA, MBC-005 was designed to provide a sustained dose of N-allyl noroxymorphone over time. We achieved this by creating a novel hydrogel, with material properties that allow it to be injectable and echogenic (i.e., observable under ultrasound). A design optimization approach was used to develop the hydrogel component of MBC-005, which resulted in elution rates for N-allyl noroxymorphone of 0.14647-mg/hr^1/2^ or <1-mg per 24-hours. This elution rate is a critical feature of any targeted therapy, such as N-allyl noroxymorphone, in which too little or too much of the drug can result in a diminished effect or, alternatively, off-target toxicity [77, 78].

In our previous work, we identified that treatment with N-allyl noroxymorphone resulted in an increase in p21 expression, which resulted in a decrease in proliferation that corresponded with SMAD1/8-dependent differentiation of mesenchymal stem cell (MSC) into osteoblasts [7]. In the current study, the BT474 cells were the most sensitive to treatment with N-allyl noroxymorphone and were observed to have an IC_50_ of 1.06-mM while the MCF7 cells were less sensitive with an IC_50_ of 2.835-mM. The normal breast epithelial cells (HUMEC) were the least sensitive to N-allyl noroxymorphone and had an IC_50_ of 4.998-mM. Given our previous work, we anticipated that cells that have a higher rate of cell division would be more sensitive to the N-allyl noroxymorphone treatment. In cultures with a higher rate of cell division we would expect that therapeutically increasing p21 expression would cause more cells to exit the cell cycle, which would lead to cell senescence and a decrease in cell number [8]. This is not what we observed in our data. In contrast, we observed that cells with a higher rate of proliferation (e.g., the MCF7 cells) were less sensitive to N-allyl noroxymorphone treatment while the BT474 cells were more sensitive. The BT474 cells were observed to have a rate of cell division that was half that of the MCF7 cells.

We also observed that treatment with N-allyl noroxymorphone resulted in an increase in p21 expression of 3.75-fold for the MCF7 cells and 5.78-fold for the BT474 cells. This increase in p21 expression is consistent with our own previous work as well as the work of others [7, 79–83]. In these studies, it was found that N-allyl noroxymorphone produced an increase in p21 expression through the opioid growth factor receptor (OGFR) [7, 79–83]. The OGFR is a receptor that binds met-5 enkephalin (Met5) that is a non-canonical homologue of the canonical opioid receptors (e.g., the μ-, κ, and δ-opioid receptors) [7, 84]. However, unlike the canonical opioid receptors, which are located on the cell surface, the OGFR is a nuclear receptor that does not bind opioids, such as morphine, but does bind morphine analogs like N-allyl noroxymorphone [7, 84]. As such, we do not believe that the increased p21 expression observed in this study and in previous work is a consequence of treatment dependent induction of cell senescence and is instead directly mediating cell exit from the cell cycle.

Putting into perspective how p21 expression coupled with the rate of cell division observed for the MCF7 and BT474 cells combined to result in the differing sensitivity to N-allyl noroxymorphone that we observed is key to understanding the mechanism of action. We believe that our results are consistent with the results presented in the study completed by Hsu et al. that posited that cell fate following exposure to doxorubicin was found to be dependent on p21 expression dynamics [9]. These authors observed that when cells were exposed to a pulse of doxorubicin that both low and high levels of p21 expression resulted in cell senescence while an intermediate level of p21 expression results in cell proliferation, which they proposed was a pathway to chemotherapy resistant cell populations. The work by Hsu et al. is also consistent with the analysis by Gewirtz, in which low dose anthracycline therapy (e.g., doxorubicin) was shown to result in cell senescence versus high doses that resulted in cell death via apoptosis [10]. In this context, we believe that the more slowly dividing BT474 cells have higher levels of endogenous p21 expression, which upon treatment with N-allyl noroxymorphone leads to an even greater increase in p21 expression levels. These elevated p21 expression level then cause the BT474 cells to exit the cell cycle, with the result of increased cell senescence. The MCF7 cells, which have a much higher rate of cell division, correspondingly would have much lower expression levels of endogenous p21 and subsequently be less sensitive to N-allyl noroxymorphone. This difference in endogenous p21 expression levels explains why it took 2x more N-allyl noroxymorphone to achieve the same level of cell senescence, as evidenced by the results of the spheroid assay.

Cells that can no longer divide are in the G_0_-phase of the cell cycle and as a result become senescent or terminally differentiate into a non-dividing cell phenotype, such as has been observed in chondrocytes or osteoblasts [7, 75, 76]. A number of authors have commented that tumor cells exiting the cell cycle and becoming senescent is associated with increased cell death [85–87]. This is consistent with the LDH results reported in this study. It is also consistent with the results for the breast epithelial cells (HUMEC), which have a reported rate of cell division that is comparable to the BT474 cells [88]. However, unlike the tumor cells, non-tumor cells, such as the HUMECs, can differentiate and exit the cell-cycle when p21 expression levels are high [8].

It is also worth noting that N-allyl noroxymorphone had inhibitory effects on cell number like those observed for treatment with doxorubicin. This effect is particularly interesting because doxorubicin is an anthracycline that is acutely toxic to both tumor cells and to non-tumor cells. In contrast, the apoptotic mechanism of cell death elicited by N-allyl noroxymorphone is only evident in tumor cells and has not been observed in non-tumor cells [7]. Further, we also identified a small but unexpected decrease in cell number and a corresponding increase in cell death from treatment with the hydrogel component of MBC-005. This effect was also specific to the MCF7 and BT474 tumor cells and was not observed in the non-tumor cells tested. This additional cytotoxic effect is intriguing; however, it is unclear how it is mediated. One possibility is the lanthanum carbonate component of the MBC-005 hydrogel. In work by Huang et al., lanthanum chloride was found to be inhibitory in two TNBC cell-lines (MDA-MB-231 and MCF10A) [89]. However, the results from the current study would appear to be the first example of a lanthanum-based therapy.

In general, all the MBC-005 doses resulted in some degree of suppressed tumor growth and increased survival. The 60-μg MBC-005 dose was the most effective, with overall survival of 33.2% and the tumor doubling time of 27-days. In addition, the Cox analysis identified that lung metastases were the event driving mortality in this model. Animals in the 60-μg MBC-005 group without lung metastases had an interpolated survival of 52.67% while the interpolated survival for animals with lung metastases in this group was 3.07%. This is in comparison to all other groups that had an interpolated survival of less than 1% when they had lung metastases.

The goal of this study was to design and test a locoregional therapeutic intervention to treat breast cancer liver metastases by optimizing the material properties of MBC-005 and then evaluating the efficacy of MBC-005 in a BALB/c mouse liver metastases model. Further, we hypothesized that an optimally designed MBC-005 would inhibit breast cancer growth *in vitro* and *in vivo*, which we have confirmed. We also hypothesized that MBC-005 would work through a p21 mechanism of action, which we have also confirmed.

Effective locoregional therapies are likely the key to reducing breast cancer mortality for patients with disseminated metastatic disease and increasing 5-year survival above 31%. In particular, localized treatments that have biological as opposed to only an ablative effect may produce more sustained and peritumoral growth arrest. Additionally, if treatment can be given when and where needed over time, the duration of overall tumor control may be enough to have favorable impacts on survival. Therapy that spares normal tissue and can be given in customized locations and appropriate dose depending on the target treatment volume could be re-administered and move disseminated metastatic breast cancer further along the spectrum toward a chronic disease.

## Conclusion

Breast cancer patients with disseminated metastatic disease that includes the liver have a poor prognosis. While a few therapies have been developed that can be delivered directly to the liver metastases, they have not shown to be effective in improving survival in breast cancer patients. However, there is a significant deficit in our knowledge due to the lack of randomized clinical trials to test these therapies. Given the heterogeneity of breast cancer phenotypes, for a randomized clinical trial to be informative it would need to be powered sufficiently to include adequate numbers of the different breast cancer phenotypes (e.g., hormone receptor types or HER2 expression levels). Intratumoral administration of MBC-005 increased survival 3.9-fold in mice and significantly decreased tumor volume 4-fold. As such, our study has demonstrated that MBC-005, a novel targeted therapy, has potential to be effective as a locoregional therapy to treat breast cancer that has metastasized to the liver in a mouse xenograft model. Further, we were able to identify an optimal dose of MBC-005. In future work, we anticipate that we will test this in phase I clinical study.

## Methods

### Material Studies

A modified USP apparatus 2 was fitted with in-situ UV fiber optic probes that was used to perform the dissolution evaluation. Solubility was assessed in 10 dissolution media (e.g., buffers with and without surfactants) to determine the ideal ‘sink’ conditions for dissolution. UV wavelengths were scanned to identify the spectra and the lambda max. The limit of detection and limit of quantification were identified, and a dissolution profile was determined. Each method was run in triplicate over 72-hours using fiber optic dip probes.

Viscosity was assessed using the SV-10 tuning-fork sine wave viscometer (A&D Weighing). Samples were prepared and mixed for 2-minutes, allowed to sit for 5-minutes, then briefly mixed a second time prior to testing. The tuning-fork sensor plates were inserted into MBC-005 and viscosity and temperature data were collected at 5-second intervals.

### In vitro Studies

Primary human mammary epithelial cells (HUMEC) were used as a non-transformed control cell line. HUMEC’s were maintained in Leibovitz’s L-15 media (Lonza), without serum but supplemented with the Mammary Epithelial Cell Growth Kit (ATCC). The MCF7 human breast cancer tumor cell-line (HR+/HER2-) is derived from a metastatic adenocarcinoma that possesses both estrogen and progesterone receptors (e.g., hormone receptor positive). The BT474 human breast cancer tumor cell-line is a ductal carcinoma derived from a solid mass (HR+/HER2+). MCF7 and BT474 cells were maintained in Dulbecco’s Modified Eagle Medium (DMEM) containing 10% fetal calf serum (FCS, v/v) and 1% penicillin–streptomycin (Cellgro, Corning Life Sciences) with 1% glutamine (Glutagro, Corning Life Sciences). All cells used in this study were obtained from ATCC.

### Cell Viability Assays

Cells were treated with serial doses of our formulation of N-allyl noroxymorphone dihydrate (Mallinckrodt Pharmaceuticals) solubilized in acetic acid, citric acid monohydrate, and 0.9% normal saline (Spectrum Chemical). Viable cell number was determined with the methyl tetrazolium (MTT) assay [75, 76, 90]. After 72-hours of treatment, 5-mg/mL of the MTT reagent (w/v, Sigma) was added to each well and incubated for 2-hours, after which the cells were lysed with 500-µL of DMSO (Sigma). 50-µL of solution was added in duplicate to a 96-well plate and MTT absorbance was measured at 570-nm on a Tecan Spark monochromator. The effects on cell proliferation were determined by normalizing treated wells relative to mean values from non-treated wells, as follows: Fold change in cell number = 100*[treated cells optical density/mean control optical density].

The CyQuant lactate dehydrogenase cytotoxicity assay (LDH; Invitrogen) was used to determine cell death following treatment with N-allyl noroxymorphone [75, 76]. Briefly, at the appropriate time points after treatment with N-allyl noroxymorphone, 50-µL of cell culture media from each experimental plate was transferred to a 96-well plate, in duplicate. 50-µL of Reaction Mixture was aliquoted into each sample well and mixed by gentle tapping. The plate was incubated for 30-minutes at 37°C in the dark, and absorbance was immediately read at 490-nm and 680-nm. The background measured at 680-nm was subtracted from the 490-nm reading to determine LDH activity. Fold change in LDH was determined using the following equation: LDH Fold Change = 100*[treated cells optical density/control well optical density].

### Spheroid Assay

The spheroid assay was used to determine long terms effects of N-allyl noroxymorphone therapy on tumor cell survival. Tumor cells in regular DMEM (5x10^4^ per implant) were mixed with equal parts (v/v) of Matrigel (Corning Life Sciences), allowed to settle in an incubator for 30-minutes, scanned using an Epson 700 scanner to determine initial spheroid area, and then carefully covered with media and placed back in the cell incubator. At the conclusion of the experiment, cells were fixed in 70% EtOH and stained with 1% crystal violet and then scanned again. Area fractions were determined using FIJI (ImageJ), and the initial area fraction for each implant was subtracted from the area fraction at the conclusion of the experiment [75].

### The p21 Assay

Cells were disassociated using a commercially available lysis buffer (Cell Signaling Technology), to which 1-mM of the protease inhibitor phenylmethylsulfonyl fluoride was added. Protein lysates were analyzed using the Pierce BCA assay to determine total protein concentrations (ThermoFisher). Total p21^Waf1/Cip1^ protein concentrations were assayed using a commercially available colorimetric sandwich ELISA assay (Cell Signaling Technology). Data were compared to a standard curve supplied with the ELISA kit.

### Mouse Xenograft Studies

The efficacy of MBC-005 in treating breast cancer metastasis in the liver was evaluated using a syngeneic mammary carcinoma model utilizing orthotopic implantation of 4T1-Luc2 cells into the intrahepatic space in female BALB/c mice (N=96 total/N=12 per group). A Cox Proportional Hazard Model was employed to assess the Hazard Ratio (HR) to determine if the following parameters contributes to animal survival: treatment (Control versus MBC-005), initial tumor volume, final tumor volume, median tumor volume, animal weight at the conclusion of the studies, metastases, bioluminescence analysis, and liver weight. A summary of the experimental design is summarized in **Figure 15**.

**Figure 15:**
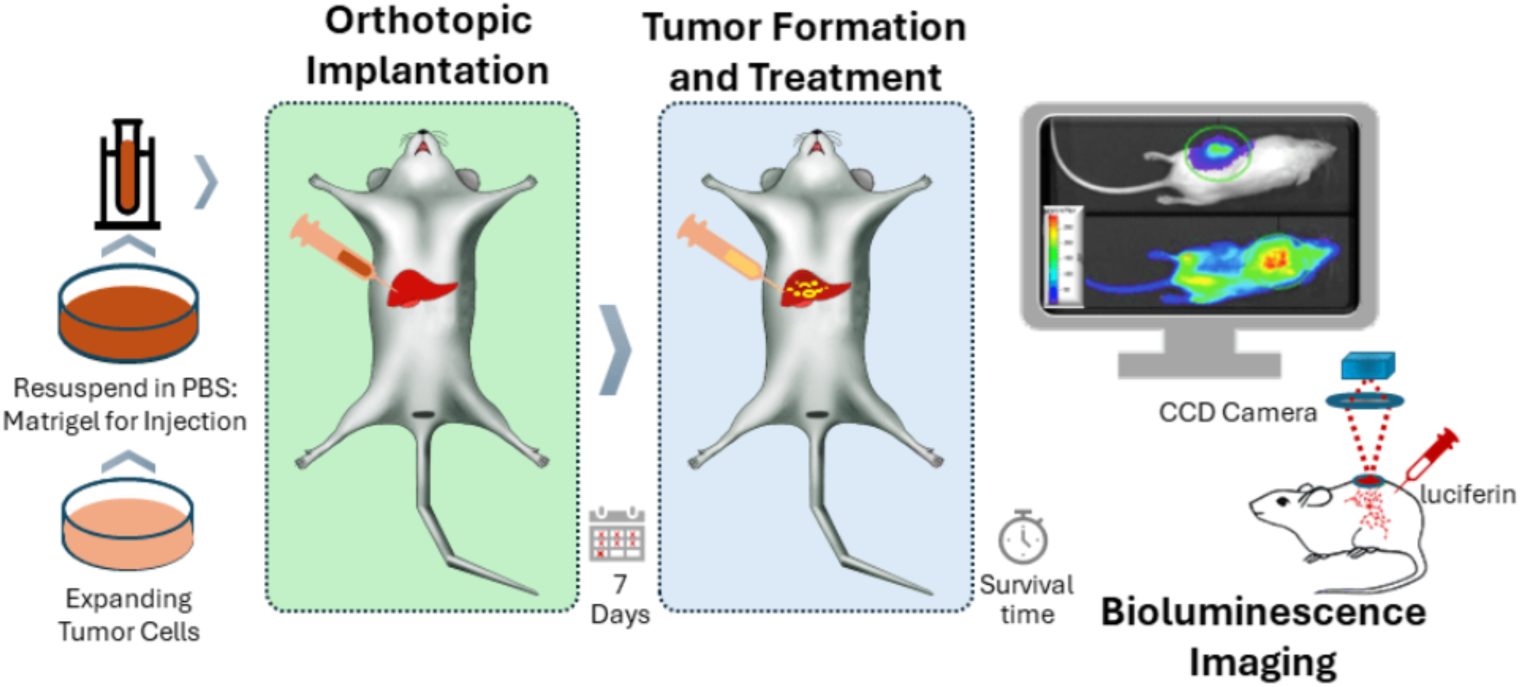
Diagrammatic illustration showing the time-course of the experimental set-up as well as the experimental endpoints.

<u>T</u>he 4T1-luc2 murine mammary gland tumor cell-line is an implantable triple-negative breast cancer that was obtained from Perkin-Elmer. The cell line was maintained in RPMI-1640 medium supplemented with 10% fetal bovine serum, 2-mM glutamine, 100-units/mL penicillin G sodium, 100-µg/mL streptomycin sulfate, and 25-µg/mL gentamicin. Cell-line bioluminescent intensity was verified by limiting dilution before and after cell implant.

For tumor implantation, the 4T1-luc2 tumor cells were resuspended in 50% Matrigel (Corning) at 4x10^6^ cells/mL.

Prior to undergoing intrahepatic implantation surgery, the animals were administered buprenorphine. Each mouse was surgically injected intra-hepatically into the left lateral lobe with 2x10^5^ cells in a 50-μL suspension under isoflurane anesthesia. On day 7 after the implantation of the 4T1-luc2 tumor cells, mice in all groups except the no treatment control group were administered with doxorubicin (5-mg/kg) intravenously. Mice in the MBC-005 treatment groups were concurrently treated with 25-μL of MBC-005 (**Table IV)** via intra-hepatically injection into the left lateral lobe.

**Table IV:** Study Design.

| Group | Number | MBC-005 Treatment |  | Doxorubicin |
| --- | --- | --- | --- | --- |
|  |  | N-allyl noroxymorphone Component | Hydrogel Component |  |
| No Treatment Control (NTC) | 12 | 0-μg | 0-μL | 0-mg/ kg |
| Control MBC-005 | 12 | 0-μg | 25-μL | 5-mg/ kg |
| 30-μg MBC-005 | 12 | 30-μg | 25-μL | 5-mg/ kg |
| 60-μg MBC-005 | 12 | 60-μg | 25-μL | 5-mg/ kg |
| 120-μg MBC-005 | 12 | 120-μg | 25-μL | 5-mg/ kg |
| 180-μg MBC-005 | 12 | 180-μg | 25-μL | 5-mg/ kg |
| 240-μg MBC-005 | 12 | 240-μg | 25-μL | 5-mg/ kg |
| 480-μg MBC-005 | 12 | 480-μg | 25-μL | 5-mg/ kg |

The surgeries, animal monitoring, and imaging were completed at Charles River Laboratory Morrisville (Morrisville, NC) in an IACUC approved study. All animal welfare considerations were taken study, including efforts to minimize suffering and distress and the use of analgesics or anesthetics. Following tumor implantation and treatment, animals were monitored daily and weighed every 2-days. Animals were euthanized immediately if they were found to be moribund or when the tumor was measured to have a caliper size greater than 1000-mm. Animals that completed the study and survived to day 60 following treatment were also euthanized. Necropsies was performed for individual animal and livers were removed for histologic processing.

### Liver Weight Measurements

Following gross necropsy, livers were collected. Liver weight was obtained before being fixed in 10% neutral buffered formalin. Liver weights were measured using a balance and corrected for the animal weight to produce a % liver weight.

### Bioluminescence Imaging

Bioluminescent images were acquired on day 0 and then bi-weekly until the end of the study. Luciferase activity was measured using the IVIS SpectrumCT (Perkin Elmer) equipped with a CCD camera, mounted on a light-tight specimen chamber. On each imaging day, animals were injected with 0.22-µm of VivoGlo D-Luciferin substrate (150-mg/kg; i.p. at 10-mL/kg, split across two injections) and placed in an anesthesia induction chamber (2.5% to 3.5% isoflurane in oxygen). Upon sedation, animals were placed in a ventral position in the imaging chamber, equipped with stage heated at physiological temperature, for image acquisition starting ten minutes after luciferin injection. Flux was quantified and reported as 10^6^ photons/second (p/s).

### Histology

After weighing the livers, samples were placed in 10% neutral buffered formalin and allowed to fix for at least 72-hours. Livers were dehydrated using serial concentrations of ethanol (e.g., 30%, 50%, 70%, and 100%) and then toluene. Livers were then laid flat, embedded in paraffin, and then sectioned at 5-μm on a rotary microtome. Sections were then stained with hematoxylin and eosin. Livers were sectioned at three levels (L1, L2, and L3). One section was evaluated from each level, which were separated by 645-μm (e.g., L1 was 645-μm from L2 and L2 was 645-μm from L3). Liver sections were imaged at 4x using a Nikon Eclipse Ni microscope and a Nikon DS-Fi2 digital camera. Composite images were generated using a tiling method, in which images were taken periodically in a grid pattern and then assembled into a final image using Photoshop (Adobe).

### Histomorphology

Histomorphometric analysis was used to determine tumor volume and the volume of necrotic tissue in the left lobe of the liver, in which the tumor and MBC-005 were implanted. Volumes for the whole liver lobe were determined using Fiji (v1.54f) using image thresholding [91]. Tumor volumes and the volume of necrotic tissue were determined using QuPath (v0.5.1-arm64) pathology analysis software [92]. Volumes were estimated based on the work of Cruz-Orive [93–95] and the Cavalieri Principle [96]. Briefly, the area fraction (*A*) for a particular level (e.g., *A_L1_*) was multiplied by the distance (*d*) between sections located in adjacent levels (i.e., between L1 and L2), in which *d*=645-μm. The composite volume (*V_t_*) for each parameter (e.g., total volume, tumor volume, of volume of necrosis) was calculated by summing the volumes for each level, such that: *V_t_* = *V_L1_* + *V_L2_* + *V_L3_* = (*A_L1_*\*645-μm) + (*A_L2_*\*645-μm) + (*A_L3_*\*5-μm). The final term *A_L3_* is multiplied by 5-μm to account for thickness of the final section. The volume fraction (*V_f_*) was calculated by dividing the tumor volume (*V_t(tumor)_*) by the total liver volume (*V_t(liver)_*). Likewise, the volume fraction of necrosis (*V_f_*) was calculated by dividing the volume of necrosis (*V_t(necrosis)_*) by the total liver volume (*V_t(liver)_*).

### Statistical Analyses

Means and standard deviations were calculated for the MTT assay data, the LDH assay data, and the spheroid assay data. The normalized dose-response data from the MTT assay were assessed using an ‘inhibitor concentration versus normalized dose response’ non-linear regression analysis. Dose-response regression lines were assessed for statistical differences between groups using an Extra-Sum-of-Squares F-test to determine if the groups were statistically different or, alternatively, could be modeled using a single regression line (α=0.05). LDH and the spheroid assay data were assessed for significant differences using either one-way ANOVA followed by correction for multiple comparisons with Dunnett’s test (α=0.05).

Bioluminescent data for the mouse studies were analyzed using a Malthusian exponential growth model was developed using non-linear least-squares regression to estimate the doubling time for the tumor volume, which served as a surrogate for local tumor growth [33]. Regression lines were assessed for statistical differences between groups using an Extra-Sum-of-Squares F-test to determine if the groups were statistically different or, alternatively, could be modeled using a single regression line (α=0.05).

Histomorphometric analyses were first evaluated normality using the Shapiro-Wilks test. All of the groups were found to be normal or lognormal, and a one-way ANOVA was used to assess differences between treatment groups. Consistent with Glantz’s et al. [97] Welch’s correction with an unpaired ‘t’ was used to identify differences between the NTC group and the MBC-005 treatment groups (α=0.05).

A Cox Proportional Hazard Regression Model [98–99] was used to assess survival and to calculate the parameter estimates (β_i_) and the hazard ratio (HR = exp(β_i_)) for the following parameters: treatment (Control versus MBC-005), initial tumor volume, final tumor volume, median tumor volume, fracture incidence, the presence of metastases, and final body weight.

All data analyses were performed using GraphPad Prism version 9.4.0, GraphPad Software, San Diego, California USA.

## Acknowledgements

Joe C. Loy for his support and editorial comments. Research reported in this publication was supported, in part, by the National Cancer Institute (NCI) of the National Institutes of Health (NIH) under awards R43CA221553 (BS Margulies) and R44CA221553 (BS Margulies).

## References

1. Choy CJ, Ling X, Geruntho JJ, Beyer SK, Latoche JD, Langton-Webster B, Anderson CJ, Berkman CE. 177Lu-Labeled Phosphoramidate-Based PSMA Inhibitors: The Effect of an Albumin Binder on Biodistribution and Therapeutic Efficacy in Prostate Tumor-Bearing Mice. Theranostics. 2017 Apr 27;7(7):1928–1939. PMID: 28638478.

2. Gangarde YM, T K S, Panigrahi NR, Mishra RK, Saraogi I. Amphiphilic Small-Molecule Assemblies to Enhance the Solubility and Stability of Hydrophobic Drugs. ACS Omega. 2020 Oct 20;5(43):28375–28381. PMID: 33163821.

3. Sun G, Rong D, Li Z, Sun G, Wu F, Li X, Cao H, Cheng Y, Tang W, Sun Y. Role of Small Molecule Targeted Compounds in Cancer: Progress, Opportunities, and Challenges. Front Cell Dev Biol. 2021 Sep 8; 9:694363. PMID: 34568317.

4. Li R, Ma XL, Gou C, Tse WKF. Editorial: Novel small molecules in targeted cancer therapy. Front Pharmacol. 2023 Aug 15;14:1272523. PMID: 37654608.

5. Mazák K, Hosztafi S, Noszál B. Species-specific lipophilicity of morphine antagonists. Eur J Pharm Sci. 2015 Oct 12;78:1–7. PMID: 26122463.

6. Mazák K, Noszál B. Lipophilicity of morphine microspecies and their contribution to the lipophilicity profile. Eur J Pharm Sci. 2012 Jan 23;45(1-2):205–10. doi: 10.1016/j.ejps.2011.11.007. Epub 2011 Nov 18. PMID: 22120645. Mazák K, Noszál B. Lipophilicity of morphine microspecies and their contribution to the lipophilicity profile. Eur J Pharm Sci. 2012 Jan 23;45(1-2):205-10. PMID: 22120645.

7. Thakur NA, DeBoyace SD, Margulies BS. Antagonism of the Met5-enkephalin-opioid growth factor receptor-signaling axis promotes MSC to differentiate into osteoblasts. J Orthop Res. 2016 Jul;34(7):1195–205. PMID: 26687326.

8. Engeland K. Cell cycle regulation: p53-p21-RB signaling. Cell Death Differ. 2022 May;29(5):946–960. PMID: 35361964.

9. Hsu CH, Altschuler SJ, Wu LF. Patterns of Early p21 Dynamics Determine Proliferation-Senescence Cell Fate after Chemotherapy. Cell. 2019 Jul 11;178(2):361–373.e12. PMID: 31204100.

10. Gewirtz DA. Does bulk damage to DNA explain the cytostatic and cytotoxic effects of topoisomerase II inhibitors? Biochem Pharmacol. 1991 Nov 27;42(12):2253–8. PMID: 1662508.

11. Rashid NS, Grible JM, Clevenger CV, Harrell JC. Breast cancer liver metastasis: current and future treatment approaches. Clin Exp Metastasis. 2021;38(3):263–277. doi:10.1007/s10585-021-10080-4

12. Zuo Q, Park NH, Lee JK, Madak Erdogan Z. Liver Metastatic Breast Cancer: Epidemiology, Dietary Interventions, and Related Metabolism. Nutrients. 2022 Jun 8;14(12):2376. PMID: 35745105.

13. Rivera K, Jeyarajah DR, Washington K. Hepatectomy, RFA, and Other Liver Directed Therapies for Treatment of Breast Cancer Liver Metastasis: A Systematic Review. Front Oncol. 2021 Mar 26;11:643383. PMID: 33842354.

14. Siegel, R. L., Miller, K. D., & Jemal, A. (2019). Cancer statistics, 2019. CA: a cancer journal for clinicians, 69(1), 7–34.

15. Siegel, R. L., Miller, K. D., & Jemal, A. (2018). Cancer statistics, 2018. CA: a cancer journal for clinicians, 68(1), 7–30.

16. DeSantis, C. E., Fedewa, S. A., Goding Sauer, A., Kramer, J. L., Smith, R. A., & Jemal, A. (2016). Breast cancer statistics, 2015: Convergence of incidence rates between black and white women. CA: a cancer journal for clinicians, 66(1), 31–42.

17. DeSantis, C. E., Ma, J., Gaudet, M. M., Newman, L. A., Miller, K. D., Goding Sauer, A., … & Siegel, R. L. (2019). Breast cancer statistics, 2019. CA: a cancer journal for clinicians, 69(6), 438–451.

18. Diamond, J. R., Finlayson, C. A., & Borges, V. F. (2009). Hepatic complications of breast cancer. The lancet oncology, 10(6), 615–621.

19. Margonis GA, Buettner S, Sasaki K, Kim Y, Ratti F, Russolillo N, Ferrero A, Berger N, Gamblin TC, Poultsides G, Tran T, Postlewait LM, Maithel S, Michaels AD, Bauer TW, Marques H, Barroso E, Aldrighetti L, Pawlik TM. The role of liver-directed surgery in patients with hepatic metastasis from primary breast cancer: a multi-institutional analysis. HPB (Oxford). 2016 Aug;18(8):700–5. doi: 10.1016/j.hpb.2016.05.014. Epub 2016 Jun 29. PMID: 27485066; PMCID: PMC4972375.

20. Liu D et al. (2020) Breast subtypes and prognosis of breast cancer patients with initial bone metastasis: a population-based study. Front Oncol 10:1. 10.3389/fonc.2020.580112.

21. Fairhurst, K., Leopardi, L., Satyadas, T., & Maddern, G. (2016). The safety and effectiveness of liver resection for breast cancer liver metastases: A systematic review. The breast, 30, 175–184.

22. Bale R, Putzer D, Schullian P. Local Treatment of Breast Cancer Liver Metastasis. Cancers (Basel). 2019;11(9):1341. Published 2019 Sep 11. doi:10.3390/cancers11091341

23. Zheng Y, Zhong G, Yu K, Lei K, Yang Q. Individualized Prediction of Survival Benefit From Locoregional Surgical Treatment for Patients With Metastatic Breast Cancer. Front Oncol. 2020 Feb 18;10:148. doi: 10.3389/fonc.2020.00148. Erratum in: Front Oncol. 2020 Apr 09;10:482. PMID: 32133290.

24. Liberchuk AN, Deipolyi AR. Hepatic Metastasis from Breast Cancer. Semin Intervent Radiol. 2020;37(5):518–526. doi:10.1055/s-0040-1720949

25. American Cancer Society, Key Statistics for Breast Cancer. Accessed 3-11 2024. https://www.cancer.org/cancer/types/breast-cancer/about/how-common-is-breast-cancer.html

26. Arnold M, Morgan E, Rumgay H, Mafra A, Singh D, Laversanne M, Vignat J, Gralow JR, Cardoso F, Siesling S, Soerjomataram I. Current and future burden of breast cancer: Global statistics for 2020 and 2040. Breast. 2022 Dec;66:15–23.

27. Li J, Xia Y, Wu Q, Zhu S, Chen C, Yang W, Wei W, Sun S. Outcomes of patients with inflammatory breast cancer by hormone receptor- and HER2-defined molecular subtypes: A population-based study from the SEER program. Oncotarget. 2017 Jul 25;8(30):49370–49379.

28. Stewart RL, Updike KL, Factor RE, Henry NL, Boucher KM, Bernard PS, Varley KE. A Multigene Assay Determines Risk of Recurrence in Patients with Triple-Negative Breast Cancer. Cancer Res. 2019 Jul 1;79(13):3466–3478.

29. Astin, Angela DiDomenico. Finger force capability: measurement and prediction using anthropometric and myoelectric measures. Diss. Virginia Tech, 1999.

30. Vo A, Doumit M, Rockwell G. The biomechanics and optimization of the needle-syringe system for injecting triamcinolone acetonide into keloids. Journal of medical engineering. 2016; 2016(1): 5162394.

31. Watt RP, Khatri H, Dibble AR. Injectability as a function of viscosity and dosing materials for subcutaneous administration. International Journal of Pharmaceutics. 2019 Jan 10;554:376–86.

32. Dai X, Cheng H, Bai Z, Li J. Breast Cancer Cell Line Classification and Its Relevance with Breast Tumor Subtyping. J Cancer. 2017 Sep 12;8(16):3131–3141. doi: 10.7150/jca.18457. PMID: 29158785; PMCID: PMC5665029.

33. Barbolosi, Dominique, et al. “Mathematical and numerical analysis for a model of growing metastatic tumors.” Mathematical biosciences 218.1 (2009): 1–14.

34. Palma D, Thakur N, Loy JC, Margulies BS. Treating bone metastases with local therapy in a breast cancer patient resulted in decreased pain and prevented fracture. Pain Manag. 2023 Oct;13(10):569–577. PMID: 37795710.

35. Adam R, Aloia T, Krissat J, Bralet MP, Paule B, Giacchetti S, Delvart V, Azoulay D, Bismuth H, Castaing D. Is liver resection justified for patients with hepatic metastases from breast cancer? Ann Surg. 2006 Dec;244(6):897–907; discussion 907-8. doi: 10.1097/01.sla.0000246847.02058.1b. PMID: 17122615; PMCID: PMC1856635.

36. Xie BJ, Zhu LN, Ma C, Li JB, Dong L, Zhu ZN, Ding T, Gu XS. A network meta-analysis on the efficacy of HER2-targeted agents in combination with taxane-containing regimens for treatment of HER2-positive metastatic breast cancer. Breast Cancer. 2020 Mar;27(2):186–196. PMID: 31529262.

37. Ji L, Cheng L, Zhu X, Gao Y, Fan L, Wang Z. Risk and prognostic factors of breast cancer with liver metastases. BMC Cancer. 2021 Mar 6;21(1):238. PMID: 33676449.

38. PDQ Treatment Information for Health Professionals: Breast Cancer Treatment (PDQ®) – Health Professional Version. Accessed 6-24 2024. https://www.cancer.gov/types/breast/hp/breast-treatment-pdq#section_7.11.

39. Harbeck, N., Penault-Llorca, F., Cortes, J. et al. Breast cancer. Nat Rev Dis Primers. 2019; 5, 66.

40. Flemming J, Madarnas Y, Franek JA: Fulvestrant for systemic therapy of locally advanced or metastatic breast cancer in postmenopausal women: a systematic review. Breast Cancer Res Treat 115 (2): 255–68, 2009.

41. Gao JJ, Cheng J, Bloomquist E, Sanchez J, Wedam SB, Singh H, Amiri-Kordestani L, Ibrahim A, Sridhara R, Goldberg KB, Theoret MR, Kluetz PG, Blumenthal GM, Pazdur R, Beaver JA, Prowell TM. CDK4/6 inhibitor treatment for patients with hormone receptor-positive, HER2-negative, advanced or metastatic breast cancer: a US Food and Drug Administration pooled analysis. Lancet Oncol. 2020 Feb;21(2):250–260.

42. Piccart M, Hortobagyi GN, Campone M, et al.: Everolimus plus exemestane for hormone-receptor-positive, human epidermal growth factor receptor-2-negative advanced breast cancer: overall survival results from BOLERO-2†. Ann Oncol 25 (12): 2357–62, 2014.

43. André F, Ciruelos E, Rubovszky G, et al.: Alpelisib for PIK3CA-Mutated, Hormone Receptor-Positive Advanced Breast Cancer. N Engl J Med 380 (20): 1929–1940, 2019.

44. Turner NC, Oliveira M, Howell SJ, Dalenc F, Cortes J, Gomez Moreno HL, Hu X, Jhaveri K, Krivorotko P, Loibl S, Morales Murillo S, Okera M, Park YH, Sohn J, Toi M, Tokunaga E, Yousef S, Zhukova L, de Bruin EC, Grinsted L, Schiavon G, Foxley A, Rugo HS; CAPItello-291 Study Group. Capivasertib in Hormone Receptor-Positive Advanced Breast Cancer. N Engl J Med. 2023 Jun 1;388(22):2058–2070. PMID: 37256976.

45. Bidard FC, Kaklamani VG, Neven P, et al.: Elacestrant (oral selective estrogen receptor degrader) Versus Standard Endocrine Therapy for Estrogen Receptor-Positive, Human Epidermal Growth Factor Receptor 2-Negative Advanced Breast Cancer: Results From the Randomized Phase III EMERALD Trial. J Clin Oncol 40 (28): 3246–3256, 2022.

46. Dowsett M, Forbes JF, Bradley R, et al.: Aromatase inhibitors versus tamoxifen in early breast cancer: patient-level meta-analysis of the randomised trials. Lancet 386 (10001): 1341–52, 2015.

47. Harris L, Batist G, Belt R, et al.: Liposome-encapsulated doxorubicin compared with conventional doxorubicin in a randomized multicenter trial as first-line therapy of metastatic breast carcinoma. Cancer 94 (1): 25–36, 2002.

48. Robert N, Leyland-Jones B, Asmar L, et al.: Randomized phase III study of trastuzumab, paclitaxel, and carboplatin compared with trastuzumab and paclitaxel in women with HER-2-overexpressing metastatic breast cancer. J Clin Oncol 24 (18): 2786–92, 2006.

49. Slamon DJ, Leyland-Jones B, Shak S, et al.: Use of chemotherapy plus a monoclonal antibody against HER2 for metastatic breast cancer that overexpresses HER2. N Engl J Med 344 (11): 783–92, 2001.

50. Sharma BB, Rai K, Blunt H, et al.: Pathogenic DPYD Variants and Treatment-Related Mortality in Patients Receiving Fluoropyrimidine Chemotherapy: A Systematic Review and Meta-Analysis. Oncologist 26 (12): 1008–1016, 2021.

51. Mamounas EP, Tang G, Fisher B, et al.: Association between the 21-gene recurrence score assay and risk of locoregional recurrence in node-negative, estrogen receptor-positive breast cancer: results from NSABP B-14 and NSABP B-20. J Clin Oncol 28 (10): 1677–83, 2010.

52. Degardin M, Bonneterre J, Hecquet B, et al.: Vinorelbine (navelbine) as a salvage treatment for advanced breast cancer. Ann Oncol 5 (5): 423–6, 1994.

53. Valero V, Forbes J, Pegram MD, et al.: Multicenter phase III randomized trial comparing docetaxel and trastuzumab with docetaxel, carboplatin, and trastuzumab as first-line chemotherapy for patients with HER2-gene-amplified metastatic breast cancer (BCIRG 007 study): two highly active therapeutic regimens. J Clin Oncol 29 (2): 149–56, 2011

54. Mathias C, Kozak VN, Magno JM, Baal SCS, Dos Santos VHA, Ribeiro EMSF, Gradia DF, Castro MAA, Carvalho de Oliveira J. PD-1/PD-L1 Inhibitors Response in Triple-Negative Breast Cancer: Can Long Noncoding RNAs Be Associated? Cancers (Basel). 2023 Sep 22;15(19):4682.

55. Howlader M, Heaton N, Rela M. Resection of liver metastases from breast cancer: towards a management guideline. Int J Surg. 2011;9(4):285–91.

56. Yoon-Flannery K, Blankenship SA, Fisher CS, Mustafa RE, Nocera NF, et al. (2018) A Systematic Review of the Surgical and Ablative Management of Breast Cancer Liver Metastasis. Adv Breast Cancer Ther: ABCT-108.

57. Timmerman RD, Bizekis CS, Pass HI, Fong Y, Dupuy DE, Dawson LA, Lu D. Local surgical, ablative, and radiation treatment of metastases. CA: a cancer journal for clinicians. 2009 May;59(3):145–70.

58. Bai X, Yang W, Zhang Z, Jiang A, Wu W, Lee J. Long-term outcomes and prognostic analysis of percutaneous radiofrequency ablation in liver metastasis from breast cancer. Int J Hyperthermia (2019) 35(1):183–93.

59. Meloni M, Andreano A, Laeseke P, Livraghi T, Sironi S, Lee F. Breast Cancer Liver Metastases: US-guided Percutaneous Radiofrequency Ablation-Intermediate and Long-term Survival Rates. Radiology (2009) 253:861–9.

60. Vogl TJ, Zangos S, Scholtz JE, Schmitt F, Paetzold S, Trojan J, Orsi F, Lotz G, Ferrucci P. Chemosaturation with percutaneous hepatic perfusions of melphalan for hepatic metastases: experience from two European centers. Rofo. 2014 Oct;186(10):937–44. Rivera et al. 13

61. Fendler WP, Lechner H, Todica A, Paprottka KJ, Paprottka PM, Jakobs TF, Michl M, Bartenstein P, Lehner S, Haug AR. Safety, Efficacy, and Prognostic Factors After Radioembolization of Hepatic Metastases from Breast Cancer: A Large Single-Center Experience in 81 Patients. J Nucl Med. 2016 Apr;57(4):517–23.

62. Liu C, Tadros G, Smith Q, Martinez L, Jeffries J, Yu Z, Yu Q. Selective internal radiation therapy of metastatic breast cancer to the liver: A meta-analysis. Front Oncol. 2022 Nov 24;12:887653.

63. Taniguchi M, Kim SR, Imoto S, Ikawa H, Ando K, Mita K, Fuki S, Sasase N, Matsuoka T, Kudo M, Hayashi Y. Long-term outcome of percutaneous ethanol injection therapy for minimum-sized hepatocellular carcinoma. World J Gastroenterol. 2008 Apr 7;14(13):1997–2002. PMID: 18395898.

64. Ebara M, Okabe S, Kita K, Sugiura N, Fukuda H, Yoshikawa M, Kondo F, Saisho H. Percutaneous ethanol injection for small hepatocellular carcinoma: therapeutic efficacy based on 20-year observation. J Hepatol. 2005 Sep;43(3):458–64. PMID: 16005538.

65. McCarley JR, Soulen MC. Percutaneous ablation of hepatic tumors. Semin Intervent Radiol. 2010 Sep;27(3):255–60. PMID: 22550364.

66. Li D, Kang J, Golas BJ, Yeung VW, Madoff DC. Minimally invasive local therapies for liver cancer. Cancer Biol Med. 2014 Dec;11(4):217–36. PMID: 25610708.

67. Ammar MB, Nouri-Neuville M, Cornelis FH. Percutaneous image-guided therapies of primary liver tumors: techniques and outcomes. La Presse Médicale. 2019 Jul 1;48(7-8): e245–50.

68. Palma DA, Olson R, Harrow S, Gaede S, Louie AV, Haasbeek C, Mulroy L, Lock M, Rodrigues GB, Yaremko BP, Schellenberg D, Ahmad B, Senthi S, Swaminath A, Kopek N, Liu M, Moore K, Currie S, Schlijper R, Bauman GS, Laba J, Qu XM, Warner A, Senan S. Stereotactic Ablative Radiotherapy for the Comprehensive Treatment of Oligometastatic Cancers: Long-Term Results of the SABR-COMET Phase II Randomized Trial. J Clin Oncol. 2020 Sep 1;38(25):2830–2838. PMID: 32484754.

69. Palma DA, Olson R, Harrow S, Gaede S, Louie AV, Haasbeek C, Mulroy L, Lock M, Rodrigues GB, Yaremko BP, Schellenberg D, Ahmad B, Griffioen G, Senthi S, Swaminath A, Kopek N, Liu M, Moore K, Currie S, Bauman GS, Warner A, Senan S. Stereotactic ablative radiotherapy versus standard of care palliative treatment in patients with oligometastatic cancers (SABR-COMET): a randomised, phase 2, open-label trial. Lancet. 2019 May 18;393(10185):2051–2058. PMID: 30982687.

70. Chalkidou A., Macmillan T., Grzeda M.T., Peacock J., Summers J., Eddy S., Coker B., Patrick H., Powell H., Berry L., et al. Stereotactic ablative body radiotherapy in patients with oligometastatic cancers: A prospective, registry-based, single-arm, observational, evaluation study. Lancet Oncol. 2021;22:98–106.

71. Tsai CJ, Yang JT, Shaverdian N, Patel J, Shepherd AF, Eng J, Guttmann D, Yeh R, Gelblum DY, Namakydoust A, Preeshagul I, Modi S, Seidman A, Traina T, Drullinsky P, Flynn J, Zhang Z, Rimner A, Gillespie EF, Gomez DR, Lee NY, Berger M, Robson ME, Reis-Filho JS, Riaz N, Rudin CM, Powell SN; CURB Study Group. Standard-of-care systemic therapy with or without stereotactic body radiotherapy in patients with oligoprogressive breast cancer or non-small-cell lung cancer (Consolidative Use of Radiotherapy to Block [CURB] oligoprogression): an open-label, randomised, controlled, phase 2 study. Lancet. 2024 Jan 13;403(10422):171–182. PMID: 38104577.

72. Tsai CJ, Yang JT, Guttmann DM, Shaverdian N, Eng J, Yeh R, Girshman J, Das J, Gelblum D, Xu AJ, Namakydoust A. Final analysis of consolidative use of radiotherapy to block (CURB) oligoprogression trial-A randomized study of stereotactic body radiotherapy for oligoprogressive metastatic lung and breast cancers. International Journal of Radiation Oncology* Biology* Physics. 2022 Dec 1;114(5):1061.

73. Tsai C.J., Yang J.T., Guttmann D.M., Shaverdian N., Shepherd A.F., Eng J., Gelblum D., Xu A.J., Namakydoust A., Iqbal A., et al. Consolidative use of radiotherapy to block (CURB) oligoprogression— Interim analysis of the first randomized study of sterotactic body radiotherapy in patients with oligoprogressive metastatic cancers of the lung and breast. Int. J. Radiat. Oncol. Biol. Phys. 2021;111:1325– 1326.

74. Chmura S.J., Winter K.A., Woodward W.A., Borges V.F., Salama J.K., Al-Hallaq H.A., Matuszak M., Milano M.T., Jaskowiak N.T., Bandos H., et al. A phase IIR/III trial of standard of care systemic therapy with or without stereotactic body radiotherapy (SBRT) and/or surgical resection (SR) for newly oligometastatic breast cancer (NCT02364557) *J*. Clin. Oncol. 2022;40:1007.

75. Margulies, B. S., Horton, J. A., Wang, Y., Damron, T. A., & Allen, M. J. (2006). Effects of radiation therapy on chondrocytes in vitro. Calcified tissue international, 78, 302–313.

76. Margulies, B. S., Damron, T. A., & Allen, M. J. (2008). The differential effects of the radioprotectant drugs amifostine and sodium selenite treatment in combination with radiation therapy on constituent bone cells, Ewing’s sarcoma of bone tumor cells, and rhabdomyosarcoma tumor cells in vitro. Journal of Orthopaedic Research, 26(11), 1512–1519.

77. Gurney H. How to calculate the dose of chemotherapy. Br J Cancer. 2002 Apr 22;86(8):1297–302. PMID: 11953888.

78. Villette CC, Orrell D, Millen J, Chassagnole C. Should personalised dosing have a role in cancer treatment? Front Oncol. 2023 May 5; 13:1154493. PMID: 37213297.

79. Cheng F, McLaughlin PJ, Verderame MF, Zagon IS. The OGF-OGFr axis utilizes the p16INK4a and p21WAF1/CIP1 pathways to restrict normal cell proliferation. Mol Biol Cell. 2009 Jan;20(1):319–27. PMID: 18923142.

80. Cheng F, McLaughlin PJ, Verderame MF, Zagon IS. The OGF-OGFr axis utilizes the p21 pathway to restrict progression of human pancreatic cancer. Mol Cancer. 2008 Jan 11;7:5. PMID: 18190706.

81. Donahue RN, McLaughlin PJ, Zagon IS. Low-dose naltrexone targets the opioid growth factor-opioid growth factor receptor pathway to inhibit cell proliferation: mechanistic evidence from a tissue culture model. Exp Biol Med (Maywood). 2011 Sep;236(9):1036–50. PMID: 21807817.

82. McLaughlin PJ, Zagon IS. The opioid growth factor-opioid growth factor receptor axis: homeostatic regulator of cell proliferation and its implications for health and disease. Biochem Pharmacol. 2012 Sep 15;84(6):746-55. PMID: 22687282.

83. Zagon IS, Porterfield NK, McLaughlin PJ. Opioid growth factor - opioid growth factor receptor axis inhibits proliferation of triple negative breast cancer. Exp Biol Med (Maywood). 2013 Jun;238(6):589–99. PMID: 23918871.

84. Zagon IS, Verderame MF, McLaughlin PJ. The biology of the opioid growth factor receptor (OGFr). Brain Res Brain Res Rev. 2002 Feb;38(3):351–76. PMID: 11890982.

85. Wang, L., Lankhorst, L. & Bernards, R. Exploiting senescence for the treatment of cancer. Nat Rev Cancer 22, 340–355 (2022).

86. Childs BG, Baker DJ, Kirkland JL, Campisi J, van Deursen JM. Senescence and apoptosis: dueling or complementary cell fates? EMBO Rep. 2014 Nov;15(11):1139–53. Epub 2014 Oct 13. PMID: 25312810.

87. Schmitt, C.A., Wang, B. & Demaria, M. Senescence and cancer — role and therapeutic opportunities. Nat Rev Clin Oncol 19, 619–636 (2022).

88. Larsson O, Blegen H, Wejde J, Zetterberg A. A cell cycle study of human mammary epithelial cells. Cell Biol Int. 1993 Jun;17(6):565–71. PMID: 8348115.

89. Huang YM, Hsu TY, Liu CY, Hsieh YC, Lai KY, Yang YW, Lo KY. Exploring the multifaceted impact of lanthanides on physiological pathways in human breast cancer cells. Toxicology. 2024 Feb; 502:153731.

90. Margulies BS, DeBoyace SD, Parsons AM, Policastro CG, Ee JS, Damron TS. Functionally deficient mesenchymal stem cells reside in the bone marrow niche with M2-macrophages and amyloid-β protein adjacent to loose total joint implants. J Orthop Res. 2015 May;33(5):615–24. PMID: 25418884.

91. Schindelin J, Arganda-Carreras I, Frise E, Kaynig V, Longair M, Pietzsch T, Preibisch S, Rueden C, Saalfeld S, Schmid B, Tinevez JY. Fiji: an open-source platform for biological-image analysis. Nature methods. 2012 Jul;9(7):676–82.

92. Bankhead P, Loughrey MB, Fernández JA, Dombrowski Y, McArt DG, Dunne PD, McQuaid S, Gray RT, Murray LJ, Coleman HG, James JA, Salto-Tellez M, Hamilton PW. QuPath: Open source software for digital pathology image analysis. Sci Rep. 2017 Dec 4;7(1):16878. doi: 10.1038/s41598-017-17204-5. PMID: 29203879; PMCID: PMC5715110.

93. Michel, R. P., & Cruz-Orive, L. M. Application of the Cavalieri principle and vertical sections method to lung: estimation of volume and pleural surface area. Journal of Microscopy. 1988; 150(2), 117–136.

94. Cruz Orive LM. On the estimation of particle number. J Microsc. 1980 Sep;120(Pt 1):15–27. doi: 10.1111/j.1365-2818.1980.tb04116.x. PMID: 7431382.

95. Cruz-Orive LM, Weibel ER (1990) Recent stereological methods for cell biology: a brief survey. Am J Physiol 258:148–156.

96. Cavalieri B. Geometria indivisibilibus continuorum: noua quadam ratione promota. Ex typographia de Ducijs. 1953.

97. Glantz SA, Slinker BK, Neilands TB. Primer of applied regression & analysis of variance, ed. McGraw-Hill, Inc., New York; 2001.

98. Spruance SL, Reid JE, Grace M, Samore M. Hazard ratio in clinical trials. Antimicrob Agents Chemother. 2004;48(8):2787–2792. doi:10.1128/AAC.48.8.2787-2792.2004

99. Barraclough H, Simms L, Govindan R. Biostatistics primer: what a clinician ought to know: hazard ratios. J Thorac Oncol. 2011 Jun;6(6):978–82.

